# A meta-analysis of sex differences in neonatal rodent ultrasonic vocalizations and the implication for the preclinical maternal immune activation model

**DOI:** 10.1101/2024.08.19.608584

**Authors:** Alison M. Randell, Stephanie Salia, Lucas F. Fowler, Toe Aung, David A. Puts, Ashlyn Swift-Gallant

## Abstract

As the earliest measure of social communication in rodents, ultrasonic vocalizations (USVs) in response to maternal separation are critical in preclinical research on neurodevelopmental disorders (NDDs). While sex differences in both USV production and behavioral outcomes are reported, many studies overlook sex as a biological variable in preclinical models of NDDs. We aimed to evaluate sex differences in USV call parameters and to determine if USVs are differently impacted based on sex in the preclinical maternal immune activation (MIA) model. Results indicate that sex differences in USVs vary with developmental stage and are more pronounced in MIA offspring. Specifically, control females exhibited longer call durations than males in early development (up to postnatal day [PND] 8), but this pattern reverses after PND8. MIA leads to a reduction in call numbers for females compared to same-sex controls in early development, with a reversal post-PND8. MIA decreased call duration and increased total call duration in males, but unlike females, developmental stage did not influence these differences. In males, MIA effects varied by species, with decreased call numbers in rats but increased call numbers in mice. The timing of MIA (gestational day ≤ 12.5 vs. >12.5) did not significantly affect the results. Our findings highlight the importance of considering sex, developmental timing, and species in USVs research. We discuss how analyzing USV call types and incorporating sex as a biological variable can enhance our understanding of neonatal ultrasonic communication and its translational value in NDD research.

## Introduction

Communication plays an essential role in survival across diverse mammalian taxa, including whales, primates, mustelids, bats, and rodents, which use high-frequency (i.e., ≤ 20 kHz) ultrasonic vocalizations (USVs) for conspecific communication (Au, 1993; Arch & Narins, 2008; Brudzynski, 2018; Clausen et al., 2008; Feng et al., 2006; Hasiniaina et al., 2018; Kruger et al., 2021; Marten, 2000; Mota-Rojas et al., 2022). Rodents are highly altricial species born with the inability to thermoregulate, for which neonates rely on maternal care (Brust et al., 2015; Ehret, 1975; Weber & Olsson, 2008). Consequently, rodent pups emit USVs, which intensify in stressful conditions, such as in periods of maternal separation (Caruso et al., 2022; Ehret, 1975; Ehret, 2005; Hammerschmidt et al., 2009). As the earliest measure of social communication and one of the first feasible behavioral tests for neonates, examining USV production in response to maternal separation is critical in preclinical research of neurodevelopmental disorders (NDDs), such as autism spectrum disorder (ASD).

Zippelius and Schleidt (1956) first classified neonate USVs as “whistles of loneliness,” expressed in response to maternal separation. Such USVs are believed to be innate signals that improve survivability by eliciting maternal retrieval (Ehret, 1975). The ventral pouch of the larynx, supported by a dorsally bent rostro-ventral component of the thyroid cartilage, contributes to the production of USVs and is developed *in utero* (Riede et al., 2017). Mouse pups emit USVs shortly after birth, although they are born deaf, with the ear canal not opening until PND10-11 (Baker et al., 2023; Ehret, 1975; Geal-Dor et al., 1993; Meng et al., 2021). Furthermore, cross-fostering studies show that mice fail to mimic USVs from other genetic strains (Kikusui et al., 2011). Together, this research suggests that pup USVs are inherent with a key role in survival.

Three types of vocalizations have been described in infant pups: 1) low-frequency (below 10 kHz) or ‘wiggling calls’ that trigger maternal licking and are produced when pups try to reach their mother’s nipple (Ehret & Bernecker, 1986); 2) broadband or ‘pain calls’ with frequencies between 4 to 40 kHz inhibit adult biting or injury and are emitted during postpartum cleaning of pups (Haack et al., 1983) and 3) isolation or distress calls (between 30 and 90 kHz) which prompt maternal retrieval and approach behaviours (Ehret, 2005). These isolation-induced USVs have increasingly received attention as a tool for assessing early communication delays in preclinical rodent models of NDDs (Scattoni et al., 2009).

### Relevance of USVs in NDDs

One of the most well-established preclinical models of ASD is the maternal immune activation (MIA) model, which consists of maternal gestational infection either via a viral mimic (e.g., polyinosinic:polycytidylic acid (poly I:C)) or bacterial agents (e.g., lipopolysaccharide (LPS)). MIA offspring show increases in ASD-like behaviors, including social communication/interaction delays and repetitive behaviors. As MIA is a risk factor for ASD in humans, and the core symptoms of ASD are recapitulated with this model, it is now one of the most studied environmental (non-genetic based) models of ASD.

Many studies that use MIA measure neonatal USV; however, there are considerable inconsistencies in both the methods and results between these studies. For instance, many report that MIA increases USV emissions (e.g., Choi et al., 2016; Kim et al., 2017), while others find decreases in MIA offspring (e.g., Carlezon et al., 2019; Malkova et al., 2012), and others report no differences (e.g., Lan et al., 2023; Straley et al., 2017). This is perhaps not surprising given the considerable variability in the analysis and reporting of isolation-induced USVs (i.e., number, duration, frequency, classifications based on call types), the species/strain of rodents used, the embryonic day (E) of MIA induction, the type of MIA (bacterial vs viral), as well as variation in the developmental stage of USV testing. Furthermore, many studies do not consider sex as a biological variable, and instead either only include males, pool the sexes, or do not report the sex of the subjects (Fig. 1A). Given the 4:1 male bias in ASD, it is critical to include sex in preclinical work to improve the translational value of findings. This is especially important in USV research, as there are significant effects of sex and sex-by-environment interactions in rodent USV production.

**Figure 1.**
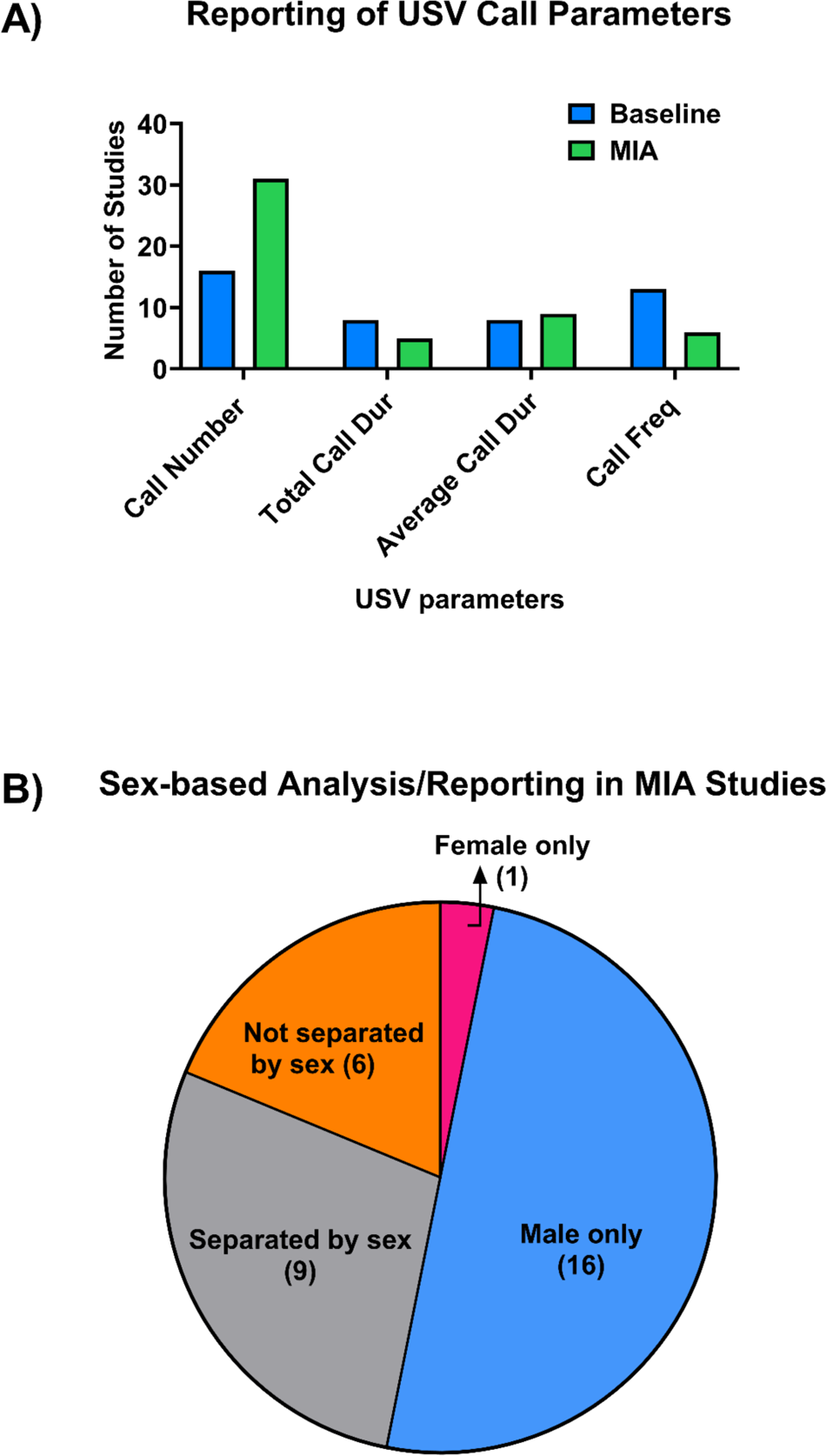
Limited USV call parameters and lack of integration of sex as a biological variable in MIA research. **A)** The most commonly reported USV call parameter is call number (49% of papers), while total call duration (10%), average call duration (17%), and call frequency (12%) are each reported in less than a fifth of studies. **B)** Less than a third of MIA studies, analyzed USVs by sex, with the majority of studies only including males or pooling the sexes in their analysis. Only 1 study analyzed USVs in females alone.

### USV Production Varies by Sex and Environmental Condition

USV production and structure differ between pup sexes, with males generally producing more USVs of longer durations, lower frequencies, and lower amplitudes than females (Caruso et al., 2020; Lenell et al., 2021). Such differences may contribute to signal saliency and subsequent maternal retrieval, as dams allocate more attention to male pups than females (Asaba et al., 2014). For example, all male litters receive more maternal care than all female litters (Alleva, 1989), and when in stressful conditions, mothers produce female-biased litters to optimize their fitness (Firman, 2020). Neuroendocrinological mechanisms, such as hypothalamic-pituitary axis (HPA) activation and co-expression of *FOXP1* and androgen receptors in the striatum, are involved in mediating sex differences in USV variation (Bowers et al., 2013; Coutellier et al., 2008; Fröhlich et al., 2017).

Pup USVs are also impacted by their rearing environment and developmental stage (Wilkin-Krug et al., 2022). For instance, pups reared in larger, environmentally enriched housing produce fewer USVs with shorter durations and lower frequencies than those in standard, under-stimulating conditions (Binder & Bordey, 2023; Wilkin-Krug et al., 2022). Moreover, pup USV production gradually rises following birth, peaks at PND8 in mice (Caruso et al., 2022; Scattoni et al., 2009) and PND10 in rats (Gulia et al., 2014), then decreases until stabilizing in puberty.

Environmental differences affect HPA activation and can modulate USV deficits in preclinical models of ASD, including the MIA model (Binder & Bordey, 2023; Conners et al., 2015; Coutellier et al., 2008; Wilkin-Krug et al., 2022; Zhao et al., 2021; Zuena et al., 2016). Thus, sex and sex-by-environment interactions influence USVs in rodents, and developmental stage, species, and strain may modulate sex differences in USVs. As such, depending on these various factors, sex differences may be overestimated, underestimated, or ignored in experimental research (Zuena et al., 2016), leading to misinterpretations of USV data and underrepresentation of sex as a mediating factor in translational rodent models of NDDs.

### Present Study Objectives

We conducted meta-analyses to assess whether 1) there are sex differences in neonatal isolation-induced USVs, 2) MIA alters neonatal isolated-induced USVs, and 3) USVs of males and females are differentially affected by MIA. Within the meta-analyses assessing these three main questions, we also assessed whether the timing of MIA, type of MIA (viral vs bacterial vs other-e.g., valproic acid), developmental stage (i.e., postnatal day of USV recordings), or species (rats vs mice) were moderators of sex and/or MIA differences in neonatal USV isolation calls.

## Methods

### Literature search

We conducted a search for studies that evaluated sex differences in neonatal USVs in response to brief maternal separation (referred herein as “baseline studies”) and those that assessed neonatal USVs in MIA preclinical models (referred herein as “MIA studies”). The search was performed using two major databases, PubMed, and Google Scholar. For baseline studies, we used keywords such as (“pup ultrasonic vocalizations” OR “neonate ultrasonic vocalizations” OR “isolation-induced ultrasonic vocalization”), combined with (“sex differences” OR “sex”). For MIA studies, additional keywords included (“maternal immune activation” OR “MIA” OR “poly ic” OR “lipopolysaccharide” OR “valproic acid” OR “animal model” OR “ASD”). This systematic review followed PRISMA protocol (Page et al., 2021; Fig. 2). The complete search strings are available in the supplementary material.

**Figure 2.**
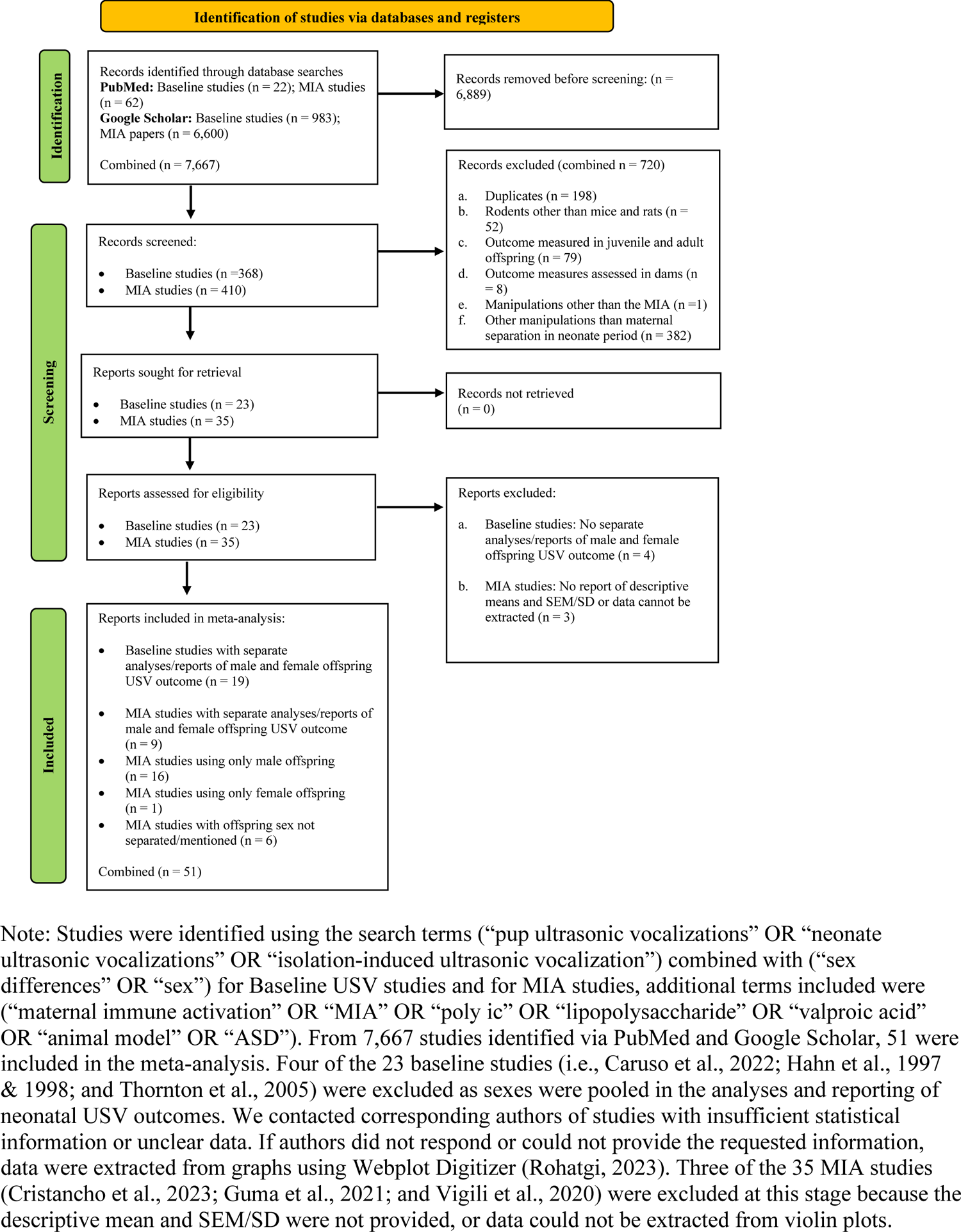
PRISMA Flowchart detailing the identification and screening of identified records for the systematic review and meta-analysis.

### Criteria for study inclusion

At the screening phase, papers were selected based on the following criteria according to the objectives of this systematic review: (i) Studies with one or more neonatal USV parameters evaluated (e.g., call number, total call duration, average call duration, and call frequency) in male and/or female offspring; (ii) USVs recorded prior to or during the weaning developmental stage (i.e., PND 3 - 21), and (iii) MIA model studies with intervention occurring during any phase of the gestational period along with appropriate controls (e.g., vehicle injection). The abstracts of all PubMed and Google Scholar records for baseline (n = 368) and MIA studies (n = 410) were evaluated for inclusion in the meta-analysis. Of 778 studies, only 23 baseline and 35 MIA independent studies included information on neonatal isolation-induced USVs recorded in rodent offspring. We excluded 4 of the 23 baseline studies (Caruso et al., 2022; Hahn et al., 1997 & 1998; and Thornton et al., 2005), as sexes were pooled in the analyses and reporting of neonatal USV outcomes.

### Extraction of study characteristics and USV parameters data

Study characteristics specific to species, strain, sex of offspring, age at which USVs were recorded, duration of USV recording, type of MIA immunogen used (e.g., poly I:C, LPS, valproic acid), dosage, gestational day of MIA induction, and frequency of administration were extracted. For USV parameters, we extracted the mean, SEM or SD, and sample size for male and female offspring (both treatment and control groups in the case of MIA studies) from each study for Hedges’s g calculation to correct for small sample bias (Turner & Bernard, 2006). We contacted corresponding authors of studies with insufficient statistical information or unclear data. If authors did not respond or could not provide the requested information, data were extracted from graphs using Webplot Digitizer (Rohatgi, 2023). Three of the 35 MIA studies (Cristancho et al., 2023; Guma et al., 2021; and Vigili et al., 2020) were excluded because the descriptive mean and SEM/SD were not provided or data could not be extracted from violin plots. When sample sizes were reported as ranges, the most conservative (i.e., lower) value was used to calculate the effect size.

### Meta-analysis

Data for the meta-analysis was analyzed using the “metafor” package in R version 4.2-0 (Viechtbauer, 2010). Some studies provided multiple measures for the same USV parameters (such as call number, mean call duration, total call duration, and call frequency) based on the developmental stage of USV recording. To address this, a three-level multilevel model was used to nest measures from the same study, correcting for the likely correlation between measures from the same study with planned subgroup analyses. First, to investigate potential sex differences in neonatal USVs in response to brief maternal separation, control samples of male and female offspring from MIA studies (n = 9 papers) were combined with the initial 19 baseline studies (total n = 28). Here, the model included species (mice vs. rats) and developmental stages (early vs. late PND) as moderators. The peak of USV production in rodents occurs around PND 8 (Scattoni et al., 2009), and as such PND 8 and below were classified as the early neonatal period, and anything above PND 8 as late. Additional moderators included, the gestational timing of MIA induction (early vs. late MIA), with gestational day (GD) 12.5 as the cut-off for early MIA and anything above GD 12.5 as late MIA, and the type of MIA immunogen used (i.e., viral injection: poly I:C vs. bacterial: LPS vs. other).

## Results

### Are there sex differences in neonatal USVs in response to brief maternal separation?

In each multilevel meta-analysis model, no significant effect size of sex differences in neonatal USVs in response to brief maternal separation was observed for call number (g = −0.01 [−0.13, 0.12], *p* = .983; SFig 1), mean call duration (g = −0.03 [−0.30, 0.25], *p* = .851; SFig 2), total call duration (g = 0.21 [−0.04, 0.47], *p* = .098; SFig 3) and call frequency (g = 0.15 [−0.11, 0.41], *p* = .259; SFig 4). We also tested whether sex differences in neonatal USVs in response to brief maternal separation were influenced by developmental stage (early vs. late PND) and species (rats vs. mice). A moderator effect was observed for mean call duration (*Q* = 6.57, *p* = .037), with a significant difference between late PND vs. early PND (g = −0.67 [−1.18, 0.16], *p* = .010; Table 1).

**Figure 3.**
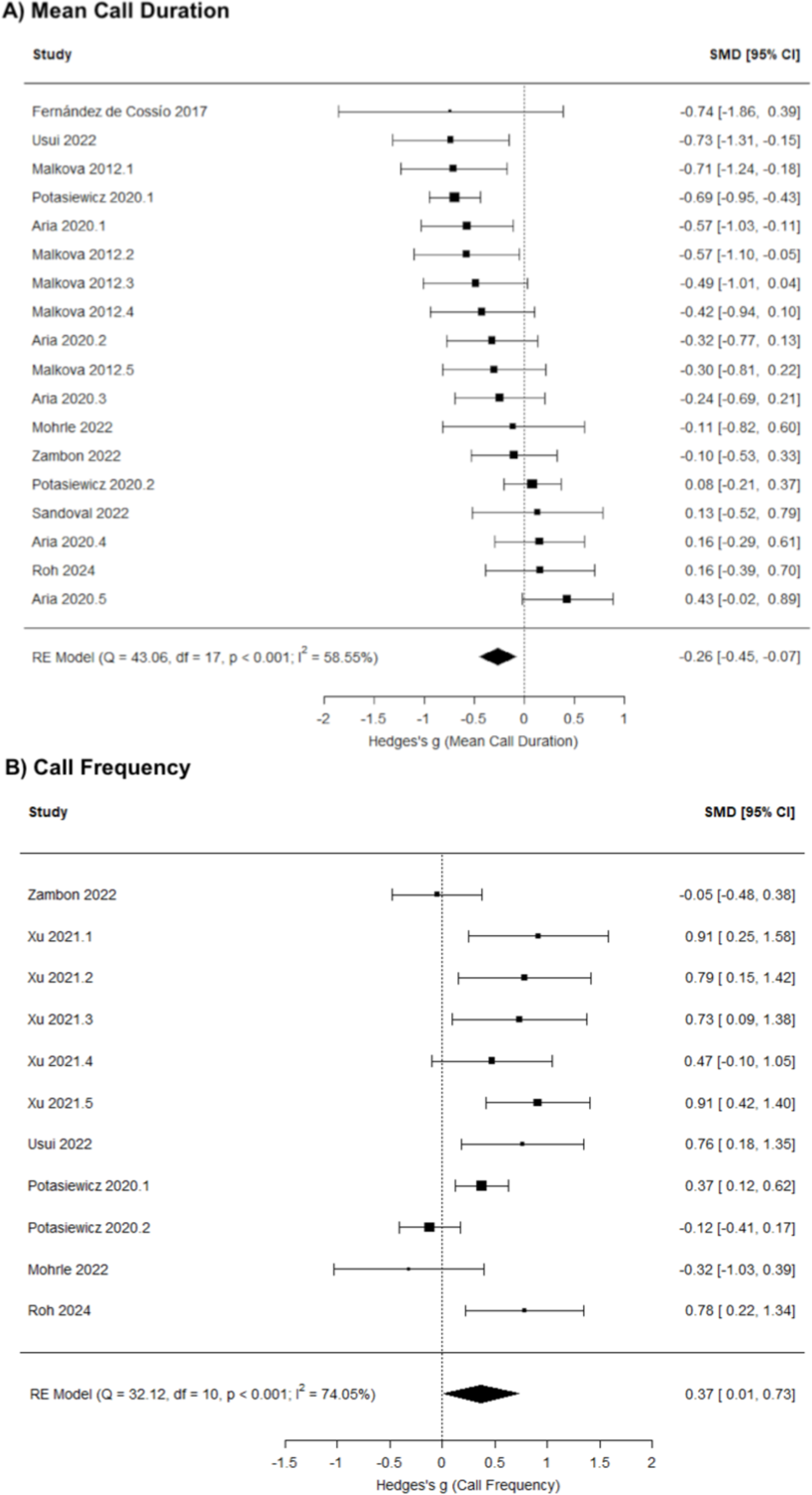
Meta-analysis results indicate that maternal immune activation (MIA) decreases neonatal USV mean call duration, and increase call frequency in response to maternal separation compared to vehicle controls

**Figure 4.**
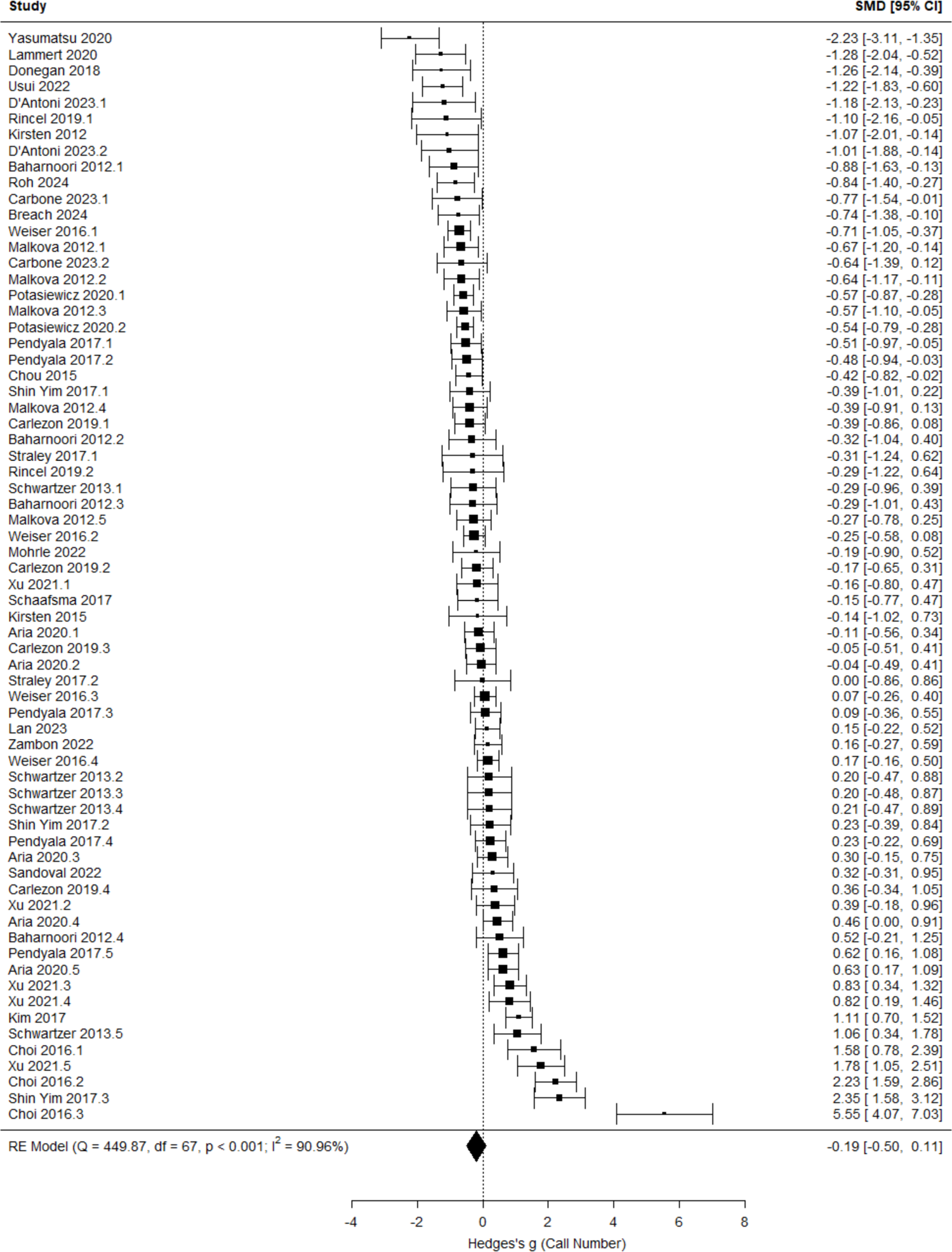
Meta-analysis results indicate that maternal immune activation does not significantly influence neonatal USV call number

**Table 1.** Results from moderator analyses for developmental stage and species.

| Models | *g* | *se* | *z* | *p* | Lower CI | Upper CI |
| --- | --- | --- | --- | --- | --- | --- |
| **Call Number** |  |  |  |  |  |  |
| Intercept | 0.06 | 0.10 | 0.61 | 0.545 | -0.14 | 0.26 |
| Late (vs. Early PND) | -0.04 | 0.12 | -0.34 | 0.731 | -0.27 | 0.19 |
| Rats (vs. Mice) | -0.18 | 0.15 | -1.18 | 0.236 | -0.48 | 0.12 |
| **Mean Call Duration** |  |  |  |  |  |  |
| Intercept | 0.28 | 0.19 | 1.47 | 0.143 | -0.09 | 0.65 |
| Late (vs. Early PND)* | -0.67 | 0.26 | -2.56 | **0.010 \*** | -1.18 | -0.16 |
| Rats (vs. Mice) | 0.16 | 0.27 | 0.59 | 0.552 | -0.37 | 0.70 |
| **Total Call Duration** |  |  |  |  |  |  |
| Intercept | 0.47 | 0.19 | 2.44 | 0.015 | 0.09 | 0.84 |
| Late (vs. Early PND) | -0.47 | 0.27 | -1.77 | 0.077 | -1.00 | 0.05 |
| Rats (vs. Mice) | -0.44 | 0.37 | -1.18 | 0.237 | -1.17 | 0.29 |
| **Call Frequency** |  |  |  |  |  |  |
| Intercept | -0.02 | 0.19 | -0.11 | 0.912 | -0.40 | 0.36 |
| Late (vs. Early PND) | 0.39 | 0.26 | 1.47 | 0.143 | -0.13 | 0.90 |
| Rats (vs. Mice) | 0.05 | 0.26 | 0.20 | 0.834 | -0.45 | 0.55 |

**Note.** \*Sub-group analyses were conducted to understand how developmental stage influences sex differences in mean call duration. Females had higher mean call duration than males in early PND groups ( $g = 0.36$ [-0.11, 0.82], $p = .130$ ), and males had higher mean call duration than females in late PND groups ( $g = -0.31$ [-0.64, 0.02], $p = .067$ ).

Sub-group analyses were conducted to understand how developmental stage influences sex differences in mean call duration. While these analyses did not reach statistical significance, the direction of the sex difference is reversed depending on developmental stage, with females having a higher mean call duration than males in early PND groups (g = 0.36 [−0.11, 0.82], *p* = .130), but males having higher mean call duration than females in late PND groups (g = −0.31 [−0.64, 0.02], *p* = .067). No other moderators were significant (*p* > .05).

### Does maternal immune activation (MIA) influence USVs in response to brief maternal separation (irrespective of sex)?

Across studies, MIA significantly influences mean call duration (g = −0.26 [−0.45, −0.07], *p* = .006; Fig. 3A) and call frequency (g = 0.37 [0.01, 0.73], *p* = .043; Fig. 3B), but not call number (g = −0.19 [−0.50, 0.11], *p* = .220; Fig. 4) or total call duration (g = 0.17 [−0.32, 0.66], *p* = .500; Fig. 5). The results of potential moderators: developmental stage (Early vs. Late PND), species (Rats vs. Mice), timing of MIA (Early vs. Late MIA), and type of MIA (Viral vs. Bacterial vs. Other) are reported in Table 2: call number (*Q* = 10.89, *p* = .054), mean call duration (*Q* = 1.99, *p* = .851), total call duration (*Q* = 18.95, *p* < .001), and call frequency (*Q* = 13.60, *p* = .009).

**Figure 5.**
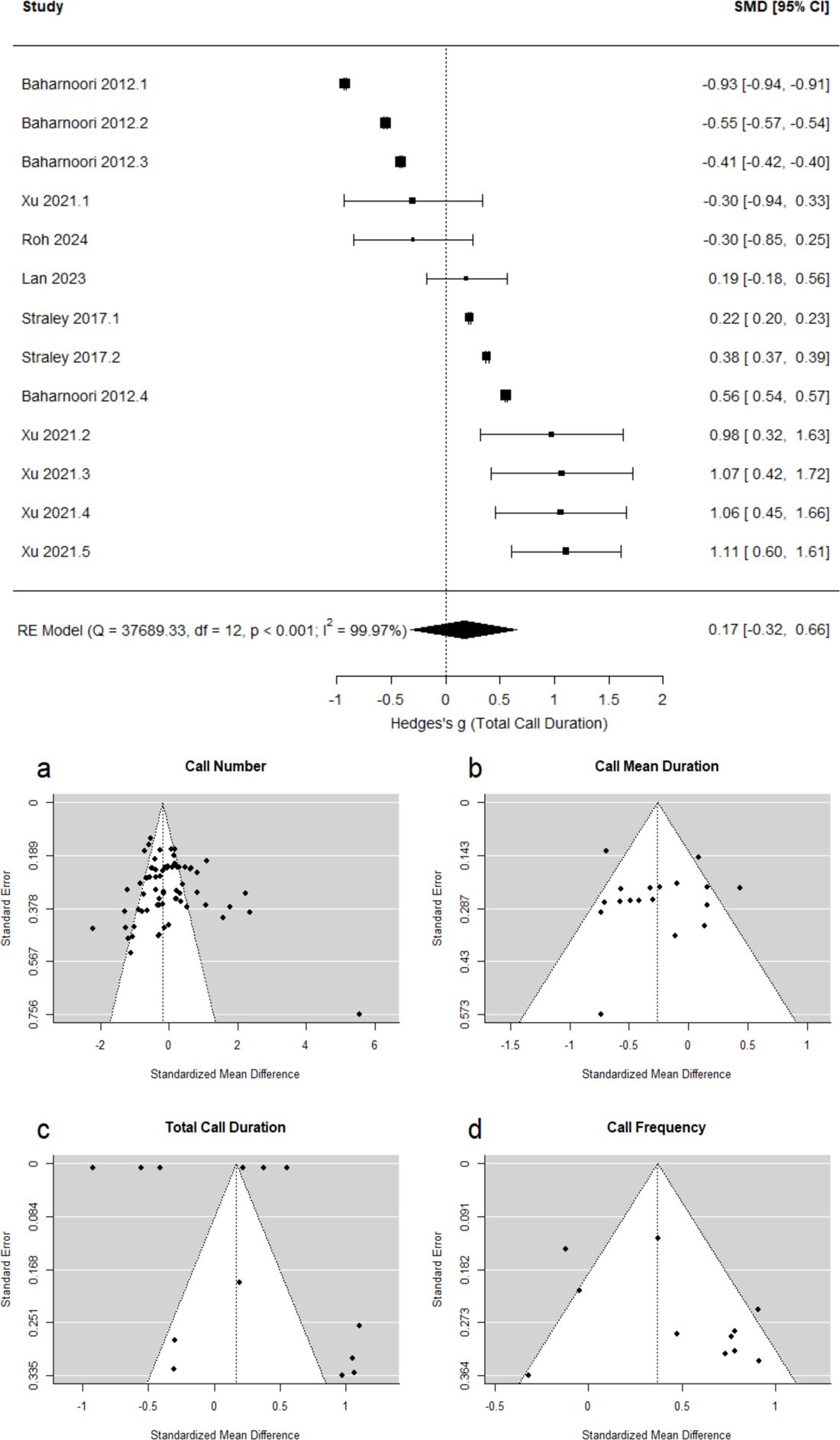
A) Meta-analysis results indicate that maternal immune activation does not significantly influence neonatal USV total call duration. B) Funnel plots of study effect sizes against standard errors for studies examining maternal immune activation on neonatal rodent USV (a) call number, (b) mean call duration, (c) total call duration, and (d) call frequency in one or both sexes.

**Table 2.**
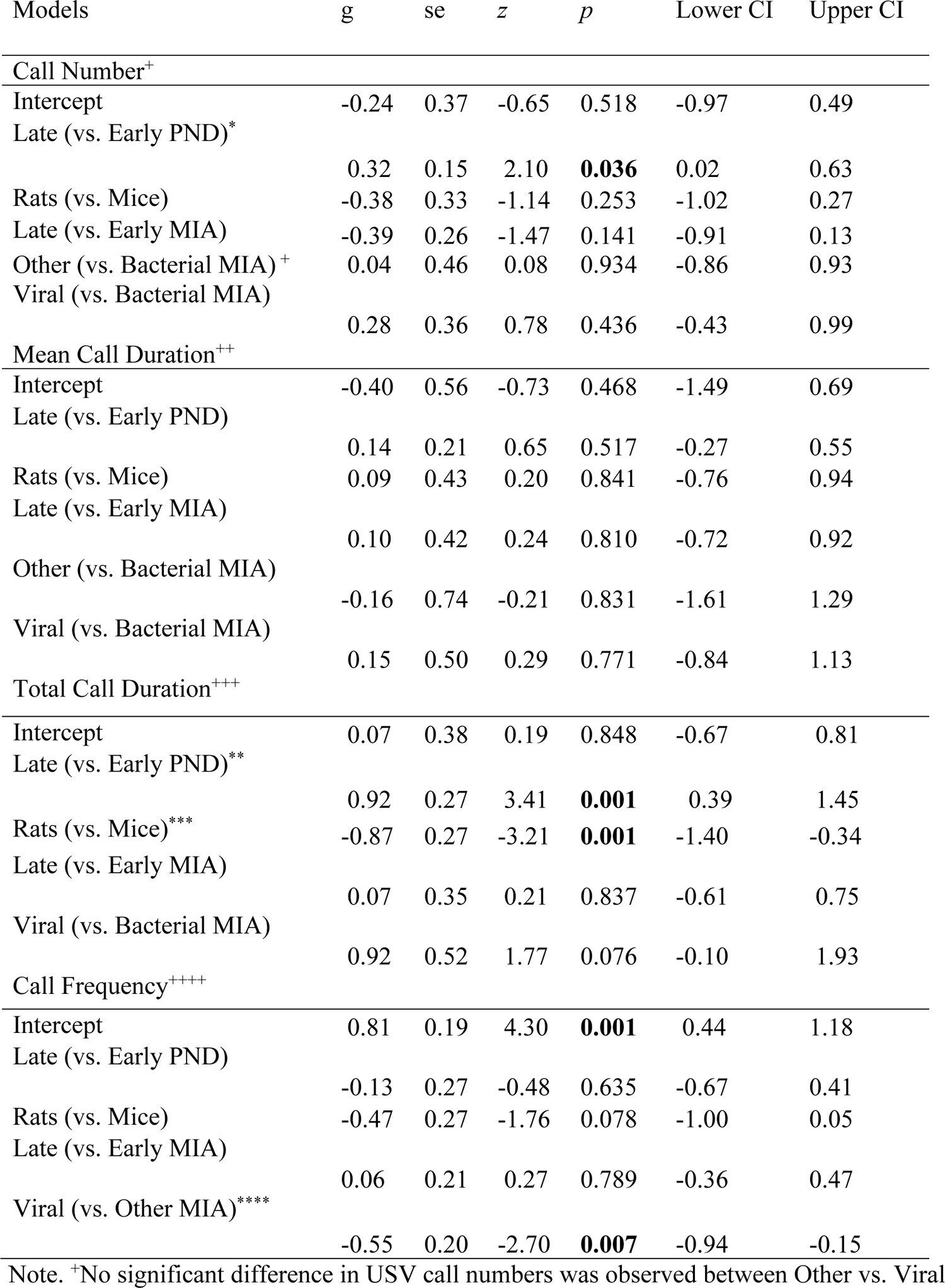
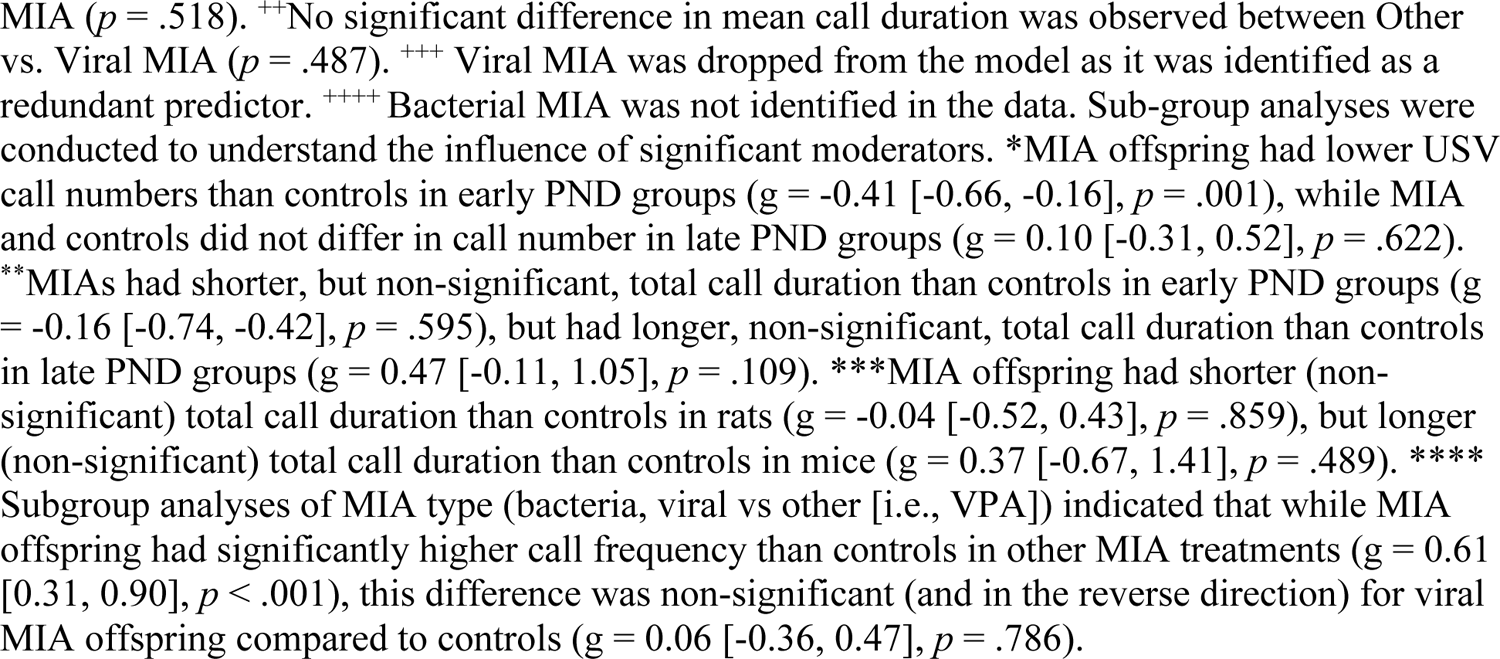
Results from moderator analyses for developmental stage, species, timing of MIA, and type of MIA for models that include one or both sexes.

| Models | g | se | z | p | Lower CI | Upper CI |
| --- | --- | --- | --- | --- | --- | --- |
| **Call Number<sup>+</sup>** |  |  |  |  |  |  |
| Intercept | -0.24 | 0.37 | -0.65 | 0.518 | -0.97 | 0.49 |
| Late (vs. Early PND)* |  |  |  |  |  |  |
|  | 0.32 | 0.15 | 2.10 | **0.036** | 0.02 | 0.63 |
| Rats (vs. Mice) | -0.38 | 0.33 | -1.14 | 0.253 | -1.02 | 0.27 |
| Late (vs. Early MIA) | -0.39 | 0.26 | -1.47 | 0.141 | -0.91 | 0.13 |
| Other (vs. Bacterial MIA)<sup>+</sup> | 0.04 | 0.46 | 0.08 | 0.934 | -0.86 | 0.93 |
| Viral (vs. Bacterial MIA) |  |  |  |  |  |  |
|  | 0.28 | 0.36 | 0.78 | 0.436 | -0.43 | 0.99 |
| **Mean Call Duration<sup>++</sup>** |  |  |  |  |  |  |
| Intercept | -0.40 | 0.56 | -0.73 | 0.468 | -1.49 | 0.69 |
| Late (vs. Early PND) |  |  |  |  |  |  |
|  | 0.14 | 0.21 | 0.65 | 0.517 | -0.27 | 0.55 |
| Rats (vs. Mice) | 0.09 | 0.43 | 0.20 | 0.841 | -0.76 | 0.94 |
| Late (vs. Early MIA) |  |  |  |  |  |  |
|  | 0.10 | 0.42 | 0.24 | 0.810 | -0.72 | 0.92 |
| Other (vs. Bacterial MIA) |  |  |  |  |  |  |
|  | -0.16 | 0.74 | -0.21 | 0.831 | -1.61 | 1.29 |
| Viral (vs. Bacterial MIA) |  |  |  |  |  |  |
|  | 0.15 | 0.50 | 0.29 | 0.771 | -0.84 | 1.13 |
| **Total Call Duration<sup>+++</sup>** |  |  |  |  |  |  |
| Intercept | 0.07 | 0.38 | 0.19 | 0.848 | -0.67 | 0.81 |
| Late (vs. Early PND)** |  |  |  |  |  |  |
|  | 0.92 | 0.27 | 3.41 | **0.001** | 0.39 | 1.45 |
| Rats (vs. Mice)*** | -0.87 | 0.27 | -3.21 | **0.001** | -1.40 | -0.34 |
| Late (vs. Early MIA) |  |  |  |  |  |  |
|  | 0.07 | 0.35 | 0.21 | 0.837 | -0.61 | 0.75 |
| Viral (vs. Bacterial MIA) |  |  |  |  |  |  |
|  | 0.92 | 0.52 | 1.77 | 0.076 | -0.10 | 1.93 |
| **Call Frequency<sup>++++</sup>** |  |  |  |  |  |  |
| Intercept | 0.81 | 0.19 | 4.30 | **0.001** | 0.44 | 1.18 |
| Late (vs. Early PND) |  |  |  |  |  |  |
|  | -0.13 | 0.27 | -0.48 | 0.635 | -0.67 | 0.41 |
| Rats (vs. Mice) | -0.47 | 0.27 | -1.76 | 0.078 | -1.00 | 0.05 |
| Late (vs. Early MIA) |  |  |  |  |  |  |
|  | 0.06 | 0.21 | 0.27 | 0.789 | -0.36 | 0.47 |
| Viral (vs. Other MIA)**** |  |  |  |  |  |  |
|  | -0.55 | 0.20 | -2.70 | **0.007** | -0.94 | -0.15 |

Note. <sup>+</sup>No significant difference in USV call numbers was observed between Other vs. Viral

Sub-group analyses were conducted to understand the influence of significant moderators (see table notes, Table 2). These results suggest that call number is reduced with MIA in early developmental stages, while MIA does not differ from controls in later development. Call duration shows a reversal with age, such that MIA offspring had shorter, but non-significant, total call duration than controls in early PND groups, but had longer, non-significant, total call duration than controls in late PND groups. Species differences suggest that MIA reduces total call duration in rats, but increases total call duration in mice. Subgroup analyses of MIA type (bacteria, viral vs other [i.e., VPA]) indicated that while MIA offspring had significantly higher call frequency than controls in other MIA treatments (g = 0.61 [0.31, 0.90], *p* < .001), this difference was non-significant (and in the reverse direction) for viral MIA offspring compared to controls (g = 0.06 [−0.36, 0.47], *p* = .786).

Potential publication bias was also evaluated via funnel plots (Fig. 5a-5d) and tested using Kendall’s rank correlations: call number (τ = −0.06, *p* = .510), mean call duration (τ = −0.20, *p* = .260), total call duration (τ = −0.08, *p* = .765), and call frequency (τ = 0.16, *p* = .542).

### Does MIA affect USVs in males and females differently?

Multilevel meta-analysis models were conducted by comparing neonatal USVs between control and MIA male rodents and by comparing between control and MIA female rodents, allowing us to assess whether MIA influences USVs more in males than females. Among male rodents, MIA significantly influences mean call duration (g = −0.41 [−0.66, −0.17], *p* = .001; Fig. 6A) and total call duration (g = 0.78 [0.25, 1.31], *p* = .004; Fig. 6B), but not call number (g = −0.27 [−0.63, 0.08], *p* = .126; Fig. 7) and call frequency (g = 0.31 [−0.34, 0.95], *p* = .349; SFig. 5). Among female rodents, MIA does not significantly influence call number (g = −0.05 [−0.36, 0.27], *p* = .762; SFig. 6), mean call duration (g = −0.23 [−0.58, 0.12], *p* = .203; SFig. 7), and call frequency (g = 0.05 [−0.28, 0.37], *p* = .766; SFig. 8).

**Figure 6.**
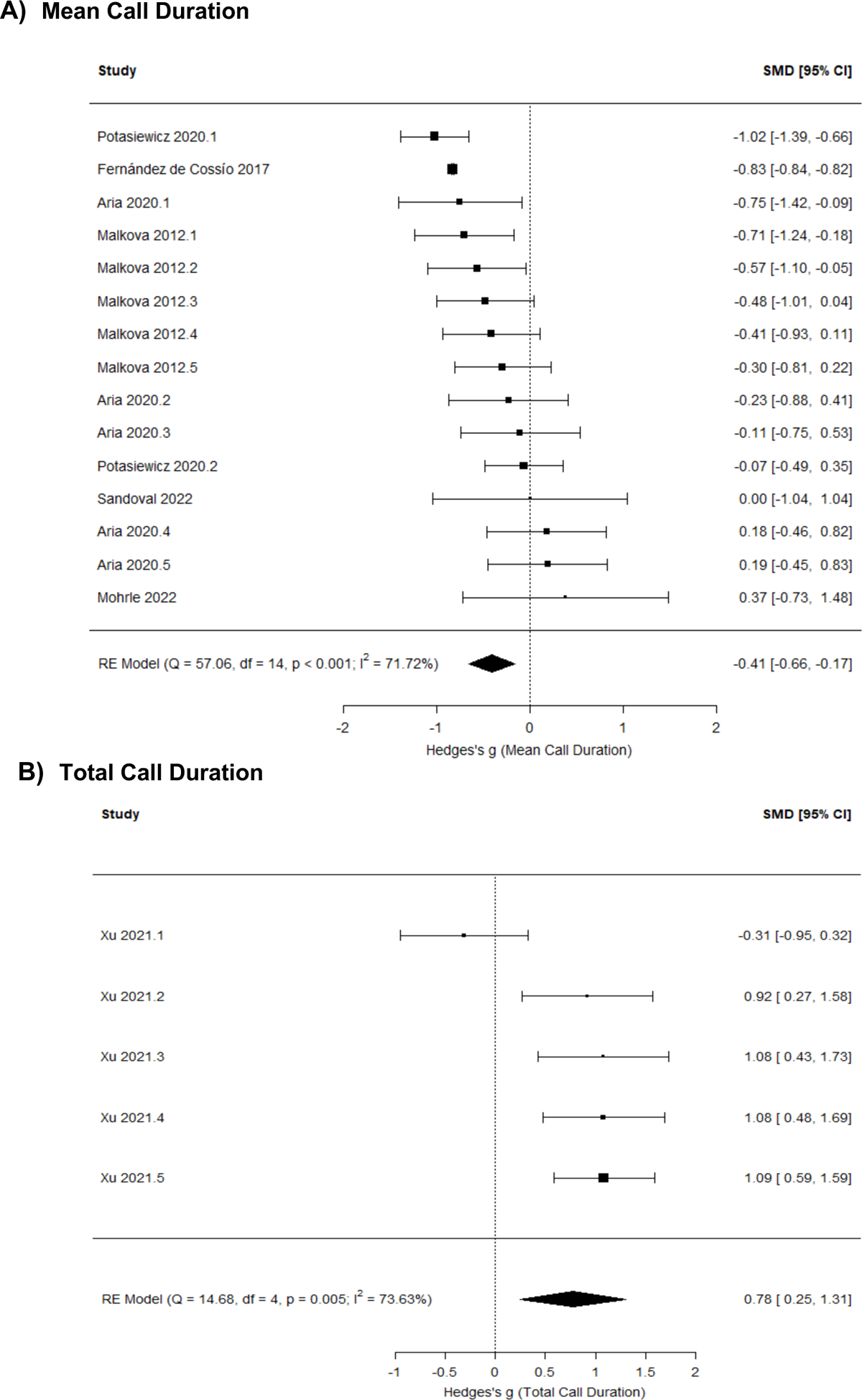
Meta-analysis results indicate that MIA decreases neonatal USV mean call duration (A) and increases total call duration (B) among male rodents compared to same-sex controls

**Figure 7.**
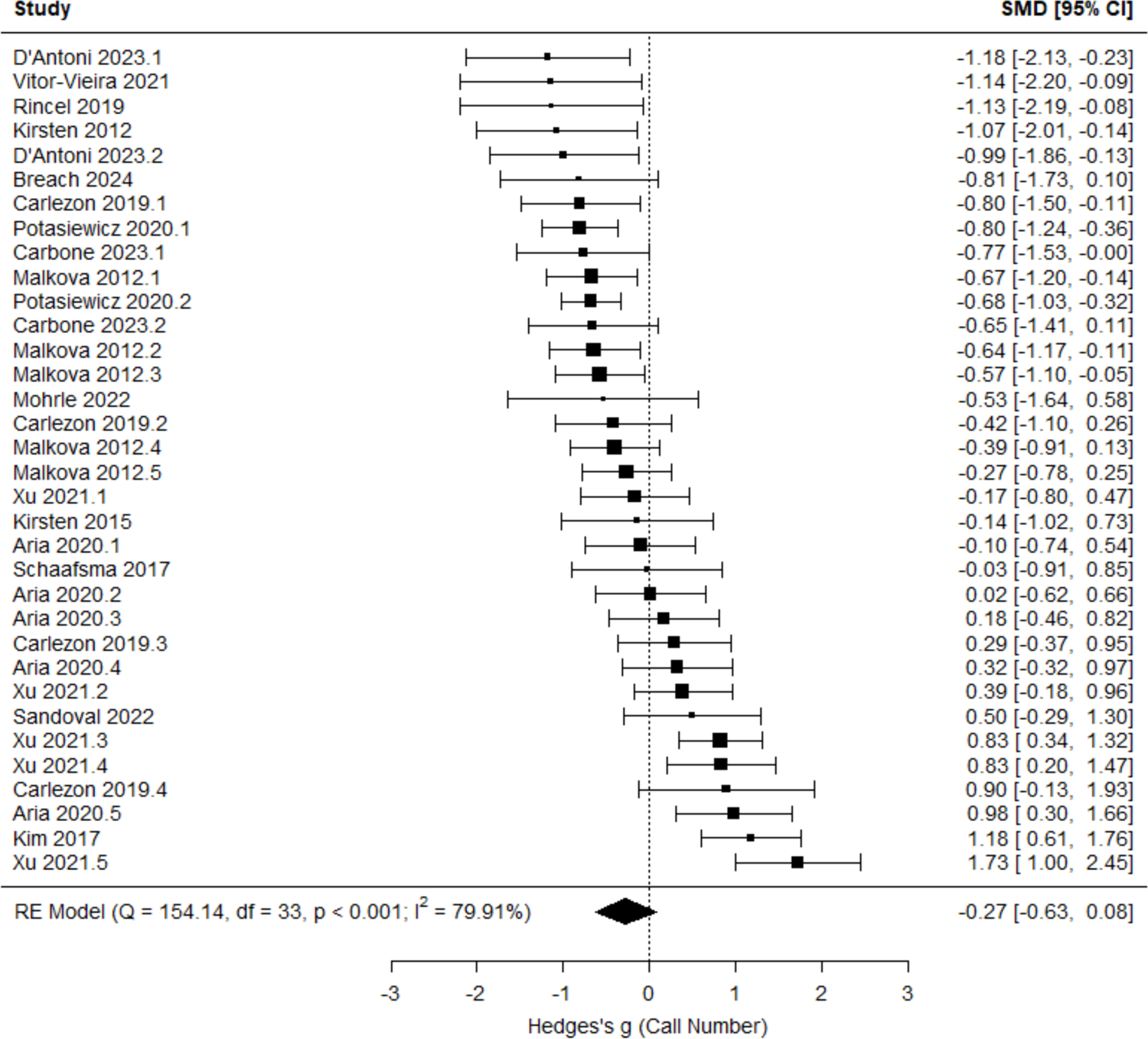
Meta-analysis results indicate that neonatal USV call number does not significantly differ between control and MIA male rodents

**Figure 8.**
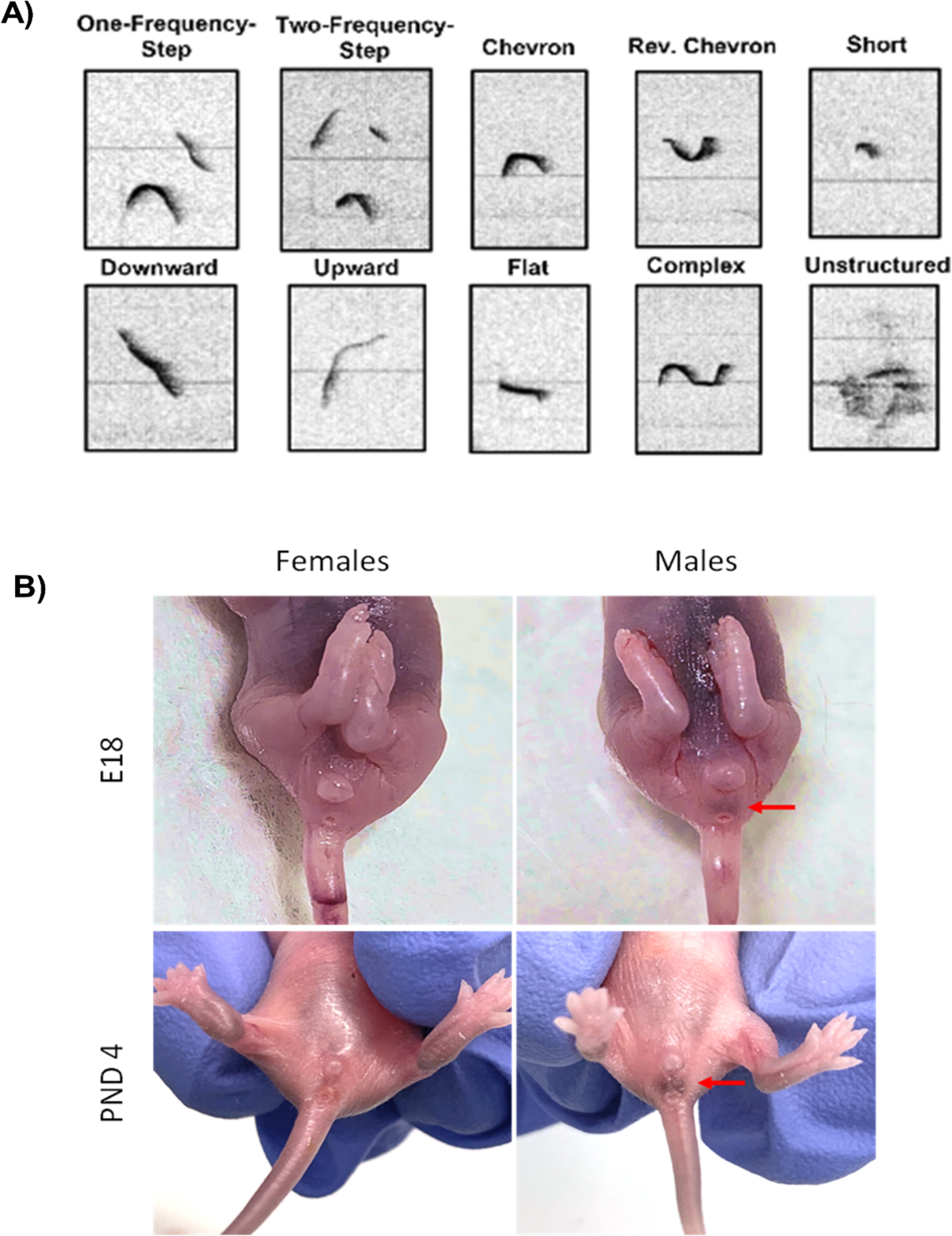
A) Call type classification (reproduced from Burke et al., 2024). B) Sexing fetal and neonatal rodents can be accomplished by visual inspection of anogenital region to identify the dark pigmented spot in the scrotal region of male pups (indicated by red arrows).

We then assessed whether sex was a significant moderator in combined analyses of male and female data. Sex was found to significantly moderate mean call duration (g = 0.06 [0.04, 0.08], *p* < .001), but not call number (g = −0.13 [−0.34, 0.09], *p* = .253) and call frequency (g = 0.19 [−0.17, 0.55], *p* = .298).

The results of potential moderators (Table 3 for males, Table 4 for females): for males, species differences were found, such that MIA reduced call numbers in rats, while in mice MIA increased call numbers compared to controls. For females, MIA reduced USV call number in early development (</=PND8), but increased call number relative to controls in later development (>PND8).

**Table 3.** Results from moderator analyses for developmental stage, species, timing of MIA, and type of MIA for models comparing control and MIA male rodents.

| Models | *g* | *se* | *z* | *p* | Lower CI | Upper CI |
| --- | --- | --- | --- | --- | --- | --- |
| **Call Numbers<sup>+</sup>** |  |  |  |  |  |  |
| Intercept | 0.13 | 0.36 | 0.35 | 0.728 | -0.58 | 0.83 |
| Late (vs. Early PND) | 0.12 | 0.20 | 0.59 | 0.557 | -0.27 | 0.50 |
| Rats (vs. Mice)* | -0.99 | 0.32 | -3.06 | **0.002** | -1.62 | -0.36 |
| Late (vs. Early MIA) | -0.20 | 0.28 | -0.70 | 0.481 | -0.76 | 0.36 |
| Other (vs. Bacterial MIA)<sup>+</sup> | 0.22 | 0.36 | 0.61 | 0.540 | -0.48 | 0.92 |
| Viral (vs. Bacterial MIA) | 0.05 | 0.34 | 0.16 | 0.873 | -0.61 | 0.72 |
| **Mean Call Duration<sup>++</sup>** |  |  |  |  |  |  |
| Intercept | -0.09 | 0.61 | -0.14 | 0.889 | -1.28 | 1.11 |
| Late (vs. Early PND) | 0.11 | 0.19 | 0.60 | 0.551 | -0.25 | 0.48 |
| Rats (vs. Mice) | 0.49 | 0.50 | 0.97 | 0.331 | -0.50 | 1.48 |
| Late (vs. Early MIA) | -0.45 | 0.55 | -0.82 | 0.414 | -1.52 | 0.62 |
| Other (vs. Bacterial MIA) | -1.43 | 0.82 | -1.74 | 0.083 | -3.04 | 0.18 |
| Viral (vs. Bacterial MIA) | -0.03 | 0.45 | -0.06 | 0.951 | -0.92 | 0.86 |
| **Total Call Duration<sup>+++</sup>** |  |  |  |  |  |  |
| Intercept | 0.38 | 0.40 | 0.95 | 0.342 | -0.40 | 1.16 |
| Late (vs. Early PND) | 0.66 | 0.51 | 1.29 | 0.196 | -0.34 | 1.65 |
| **Call Frequency<sup>++++</sup>** |  |  |  |  |  |  |
| Intercept | 0.29 | 0.66 | 0.44 | 0.662 | -1.00 | 1.57 |
| Late (vs. Early PND) | -0.13 | 0.29 | -0.45 | 0.654 | -0.69 | 0.43 |
| Rats (vs. Mice) | 0.24 | 0.68 | 0.35 | 0.724 | -1.10 | 1.58 |
| Late (vs. Early MIA) | 0.58 | 0.61 | 0.94 | 0.346 | -0.62 | 1.78 |
| Viral (vs. Other MIA) | -1.17 | 0.60 | -1.95 | 0.051 | -2.35 | 0.00 |

Note. <sup>+</sup>No significant difference in USV call numbers was observed between Other vs. Viral MIA ( $p = .623$ ). <sup>++</sup>A significant difference in mean call duration was observed between Other vs. Viral MIA ( $g = 1.40$ ; $p = .015$ ). <sup>+++</sup> Only 1 level of species, timing of MIA, and type of MIA were identified in the data, and predictors were not included in the model. <sup>++++</sup> Bacterial MIA was not identified in the data. Call number ( $Q = 11.59$ , $p = .041$ ), mean call duration ( $Q = 13.80$ , $p = .017$ ), total call duration ( $Q = 1.67$ , $p = .196$ ), and call frequency ( $Q = 15.12$ , $p = .005$ ). Sub-group analyses were conducted to understand how species influences USV call numbers in MIA male rodents. MIA male rodents had lower USV call numbers than control males in rats ( $g = -0.74$ [-0.95, -0.53], $p < .001$ ), and had higher USV call numbers than control males in mice ( $g = 0.14$ [-0.30, 0.58], $p = .534$ ).

**Table 4.** Results from moderator analyses for developmental stage, species, timing of MIA, and type of MIA for models comparing control and MIA female rodents.

| Models | *g* | *se* | *z* | *p* | Lower CI | Upper CI |
| --- | --- | --- | --- | --- | --- | --- |
| **Call Numbers<sup>+</sup>** |  |  |  |  |  |  |
| Intercept | -0.15 | 0.37 | -0.40 | 0.687 | -0.87 | 0.57 |
| Late (vs. Early PND)* | 0.58 | 0.28 | 2.03 | **0.043** | 0.02 | 1.13 |
| Rats (vs. Mice) | 0.05 | 0.49 | 0.10 | 0.917 | -0.90 | 1.01 |
| Late (vs. Early MIA) | -0.07 | 0.35 | -0.21 | 0.832 | -0.76 | 0.61 |
| Other (vs. Bacterial MIA)<sup>+</sup> | -0.27 | 0.64 | -0.42 | 0.674 | -1.53 | 0.99 |
| Viral (vs. Bacterial MIA) | -0.13 | 0.43 | -0.31 | 0.758 | -0.99 | 0.72 |
| **Mean Call Duration<sup>++</sup>** |  |  |  |  |  |  |
| Intercept | -1.09 | 1.00 | -1.09 | 0.275 | -3.05 | 0.87 |
| Late (vs. Early PND) | 0.42 | 0.42 | 1.01 | 0.315 | -0.40 | 1.24 |
| Rats (vs. Mice) | 0.05 | 1.14 | 0.04 | 0.967 | -2.19 | 2.29 |
| Late (vs. Early MIA) | 0.60 | 0.93 | 0.65 | 0.518 | -1.22 | 2.42 |
| Other (vs. Bacterial MIA) | 0.60 | 1.99 | 0.30 | 0.764 | -3.31 | 4.51 |
| Viral (vs. Bacterial MIA) | 0.67 | 1.31 | 0.52 | 0.606 | -1.89 | 3.23 |
| **Call Frequency<sup>+++</sup>** |  |  |  |  |  |  |
| Intercept | 0.26 | 0.19 | 1.39 | 0.164 | -0.11 | 0.62 |
| Late (vs. Early MIA) | -0.01 | 0.52 | -0.03 | 0.977 | -1.03 | 1.00 |
| Viral (vs. Other MIA) | -0.38 | 0.51 | -0.75 | 0.454 | -1.38 | 0.62 |

Note. Total call duration data were not identified for analysis. <sup>+</sup>No significant difference in USV call numbers was observed between Other vs. Viral MIA ( $p = .713$ ). <sup>++</sup>No significant difference in mean call duration was observed between Other vs. Viral MIA ( $p = .936$ ). <sup>+++</sup>Only 1 level of developmental stage and species were identified in the data and were not included in the model. Bacterial MIA was not identified in the data. \*Sub-group analyses were conducted to understand how developmental stage influences USV call numbers in MIA female rodents. MIA female rodents had lower USV call numbers than control females in early PND groups ( $g = -0.25$ [-0.49, -0.01], $p = .038$ ), and had higher USV call numbers than control females in late PND groups ( $g = 0.36$ [-0.21, 0.93], $p = .217$ ). Call number ( $Q = 6.51$ , $p = .260$ ), mean call duration ( $Q = 2.88$ , $p = .718$ ) and call frequency ( $Q = 2.25$ , $p = .325$ ).

## Discussion

The present meta-analyses revealed that developmental stage is a significant moderator of sex differences in USVs, and sex differences in USVs are more pronounced in the preclinical MIA model. While USVs differed between MIA and control offspring when data were pooled across sexes, these differences should be interpreted with caution: Analyses by sex revealed that sex differences in USVs depend on developmental stage and species. Moreover, the majority of differences were found in mean or total call duration; this is an important consideration for future research, as many studies evaluating early communication delays via USVs report only call number (see Fig. 1B), without any measure of call duration. The differences in mean call duration, but not always in total call duration or call number, may also suggest that there are group differences in call types (i.e, flat, two-step, chevron, etc., see Fig. 8A). As such, we recommend that in studying USVs in MIA (or other preclinical models), researchers consider developmental stage, species, sex, and expand their analysis to include additional USV parameters, including call type. Below, we detail the significance of these findings, as well as provide recommendations for non-invasive visual sexing of neonates, and for automated scoring software that allows for accurate and relatively quick analyses of USV call parameters and call type classification.

We found that at baseline, males and females differ in USVs depending on developmental stage. Specifically, developmental stage was a significant moderator of sex differences in USVs, and trending subgroup analyses indicate that females display longer mean call durations than males early in development (≤ PND8), but this reverses later (> PND8), with males calling longer. This aligns with known sex differences in maternal care. Specifically, throughout development male pups generally receive greater maternal investment, as indicated by more anogenital licking and nursing, lower pup-retrieval latencies, and more maternal aggression towards intruders relative to females (Alleva et al., 1989; Musi et al., 1993; Moore & Morelli, 1979). However, such differences are more pronounced later in development; for instance, Long Evans Rats show marginal sex differences in maternal anogenital licking at PND3 but stark differences on PND7, 10, and 14 (Moore & Morelli, 1979). Similarly, sex differences in anogenital licking are lower at PND2 than PND10 in House Mice (Moore, 1992). One possibility is that females may call longer at early development because sex differences in maternal behaviour are still marginal, meaning they may be calling more in an attempt to outcompete male littermates (Ehret, 2005). However, at later PNDs, female pups may be experiencing learned helplessness in their USV production, where they call less in response to being outcompeted by males for maternal attention. On the other hand, males may call more in later development as maternal care is required for male-specific development and socio-sexual behaviours (e.g., courtship; Rilling & Young, 2014). Regardless, the present meta-analysis suggests that researchers should consider developmental timing when studying USVs generally and in translational NDD models, as PND can modulate sex differences in USV emission and, potentially, maternal response.

When assessing the effects of MIA across all studies (i.e., pooling sexes), MIA reduced mean call duration and increased call frequency, and when developmental stage is considered, call number reductions were found to be restricted to early development. Species was also a significant moderator, such that call duration is reduced in MIA rats but increased in MIA mice. However, when sex is considered as a biological variable, it can be discerned that many of these effects are driven by males *or* females. For example, MIA decreased mean call duration, but increased total call duration in males, but not in females. However, when developmental stage is considered for females, MIA reduced USV call numbers in early development (≤ PND8) relative to controls, but increased in later development (> PND8). Moreover, species differences were found only for males, such that MIA reduced call numbers in rats, while in mice MIA increased call numbers compared to controls. These findings highlight that USVs are affected by MIA in both males and females, but are moderated by developmental stage and species.

Of note, sex differences in the presentation of USV delays underscores the necessity of considering sex as a biological variable in USV and NDD research, especially in light of the sex difference in the clinical manifestations of these conditions. For instance, in ASD, there is a known gender bias in diagnosis, where females who meet the criteria of ASD are less likely to receive a clinical diagnosis (Loomes et al., 2017) or receive a diagnosis later in life (Begeer et al., 2013; Taylor et al., 2019). This diagnostic disparity may partly result from variations in symptom presentation. This bias extends into preclinical research, where studies frequently focus on males or fail to account for sex differences, leading to incomplete results and skewed conclusions (e.g., Lammert & Lukens, 2019; Lan et al., 2023; Malkova et al., 2012; Straley et al., 2017). The present study highlights that neonatal USV production in response to MIA is affected in both males and females, but additional research is necessary to gain a comprehensive understanding of the effects of MIA in the understudied sex - females.

Indeed, most studies in the present meta-analysis were single-sex (17 of 32), and 40% of studies reporting both sexes pooled male and female data (6 of 15 studies) for assessing USVs; although, we note that pooling of the sexes was not always the case for other juvenile behavioral measures in these studies. We hypothesize this may be due to unfamiliarity in sexing neonatal pups or the invasive nature of genotyping and/or tracking into adolescence. However, accurate sexing of neonates, and even fetuses at embryonic day 18, via visual inspection alone is feasible. Our data show fetal sexing accuracy is > 93% (n = 39 of 43), and sexing from PND 4 is > 97% accurate (n = 481 of 492), in C57BL/6 mice. As illustrated in Fig. 8B, males and females show distinct anogenital differences by late embryonic development: males have a noticeable dark spot in the scrotal region, which is absent in females. This pigmentation is the result of higher melanin concentration linked to androgen exposure (Wilson & Spaziani, 1973). It is visible from birth in several dark-fur mouse strains (Wolterink-Donselaar et al., 2009) and Long-Evans rats (Wilson & Spaziani, 1973), although white-furred strains lack this pigmentation (e.g., A/J, 129X1/SvJ, and C57Bl/6J-Chr 7A/NaJ; Wolterink-Donselaar et al.). Thus, we recommend visual inspection for sexing in dark-furred rodent strains, without genotyping and/or tattooing for tracking, and urge reviewers and editors to accept this method as valid, to promote the inclusion of sex as a biological variable in neonatal rodent studies.

In addition to including sex, expanding USV analysis to include the wide repertoire of USV call types emitted by pups may provide insight into the characteristics of pups’ ultrasounds associated with NDD-like phenotypes. Indeed, USV call types, categories or classifications, since their description by Sales and Smith (1978), have become an integral part of qualitative analysis of rodent vocalizations. These spectrographic analyses are based on the frequency modulation and duration of acoustic signals (Branchi et al., 1998) and may provide additional information for quantitative USV parameters, such as call number, duration, and frequency (Branchi et al., 1998; Caruso et al., 2022; Scattoni et al., 2008). Before Scattoni et al. (2008), USVs were classified into five categories: constant frequency, modulated frequency, frequency steps, composite, and short (Branchi et al., 1998; Sales & Smith, 1978). Scattoni’s taxonomy expanded this framework to include additional call types—complex, harmonics, two-syllable, upward, downward, flat, chevron, short, composite, and frequency steps (Fig. 8A)—allowing for the detection of more subtle differences (Caruso et al., 2022; Scattoni et al., 2008). Functionally, specific USV call types have been suggested to correlate with later social behavior. For example, infant isolation-induced calls, such as flat and short calls, have been found to correlate with the frequency of social interactions during adolescence (Granata et al., 2022). This indicates that analyzing USV call type repertoire in pups can be an additional tool for quantifying the extent of ASD-like traits in rodent models during early development (Möhrle et al., 2023).

Several automated software programs (e.g., Avisoft analysis, VocalMat, DeepSqueak, Kaleidoscope Pro) can automatically segment rodent audio files into calls and apply classification algorithms to label vocalizations (for detailed reviews see Coffey et al., 2019; Fonseca et al., 2021 and Premoli et al., 2021). Our lab currently uses Kaleidoscope Pro, in which users can use advanced classifiers to recognize a predetermined number of call types. This allows us to accurately sort and classify various calls, significantly accelerating the analysis of USVs. We have successfully classified over 70,000 USV calls with an accuracy rate exceeding 80%, which we can then manually correct to reach near 100% accuracy. We are able to share these training data with interested researchers upon request.

In conclusion, our meta-analyses reveal notable sex differences in USVs, both at baseline and in response to MIA. However, these differences vary depending on developmental stage and species. Importantly, none of the analyses showed differences across all USV call parameters, indicating a need for researchers to expand their focus beyond call number to include at minimum call duration and mean call duration. We also recommend incorporating an analysis of USV call types, as the existing studies, while few at this point, have identified sex-specific differences in call types (Granata et al., 2021; Kentner et al., 2018). Additionally, we highlight that visual inspection via the “spot method” (Wolterink-Donselaar et al., 2009) is a highly accurate method for determining sex in common laboratory strains (e.g., C57Bl/6 mice and Long Evans rats) and advocate for its use to encourage the inclusion of sex in neonatal research.

## Supplementary Data

### Baseline paper search terms

#### Google Scholar

“pup ultrasonic vocalizations”|”neonate ultrasonic vocalization”|”maternal separation usv”|”isolation-induced ultrasonic vocalization”and “sex differences”|”sex”|”males”|”females”|”animal experimentation” ==983 until June 2024

#### PubMed

”pup ultrasonic vocalization*” OR “isolation-induced ultrasonic vocalization*” OR “neonate ultrasonic vocalization*” OR “infant ultrasonic vocalization*” OR “maternal separation ultrasonic vocalization*” OR “neonate USV*” OR “pup USVs*” AND “sex differences*” == 22 papers

### MIA papers search terms

#### Google Scholar

”maternal immune activation”|”MIA”|”poly ic”|”poly i:c”|”poly(i:c)”|”Polyinosinic:polycytidylic acid”|”lipopolysaccharide”|”lps”|”autism”|”autism spectrum disorder”|”asd”|”valproic acid” and “isolation-induced ultrasonic vocalizations”|”ultrasonic vocalization”|”usv”|”usvs”| and “sex”|”sex differences”|”male”|”males”|”female”|”females”|”animal experimentation” = 6600

#### PubMed

“maternal immune activation” OR “Polyinosinic:polycytidylic acid” OR “poly ic” OR “poly i:c” OR “lipopolysaccharide” OR “lps” “valproic acid” AND “isolation-induced ultrasonic vocalizations” OR “neonate ultrasonic vocalization” OR “pup usvs” AND “sex” = 62

---

**Supplemental Figure 1.**
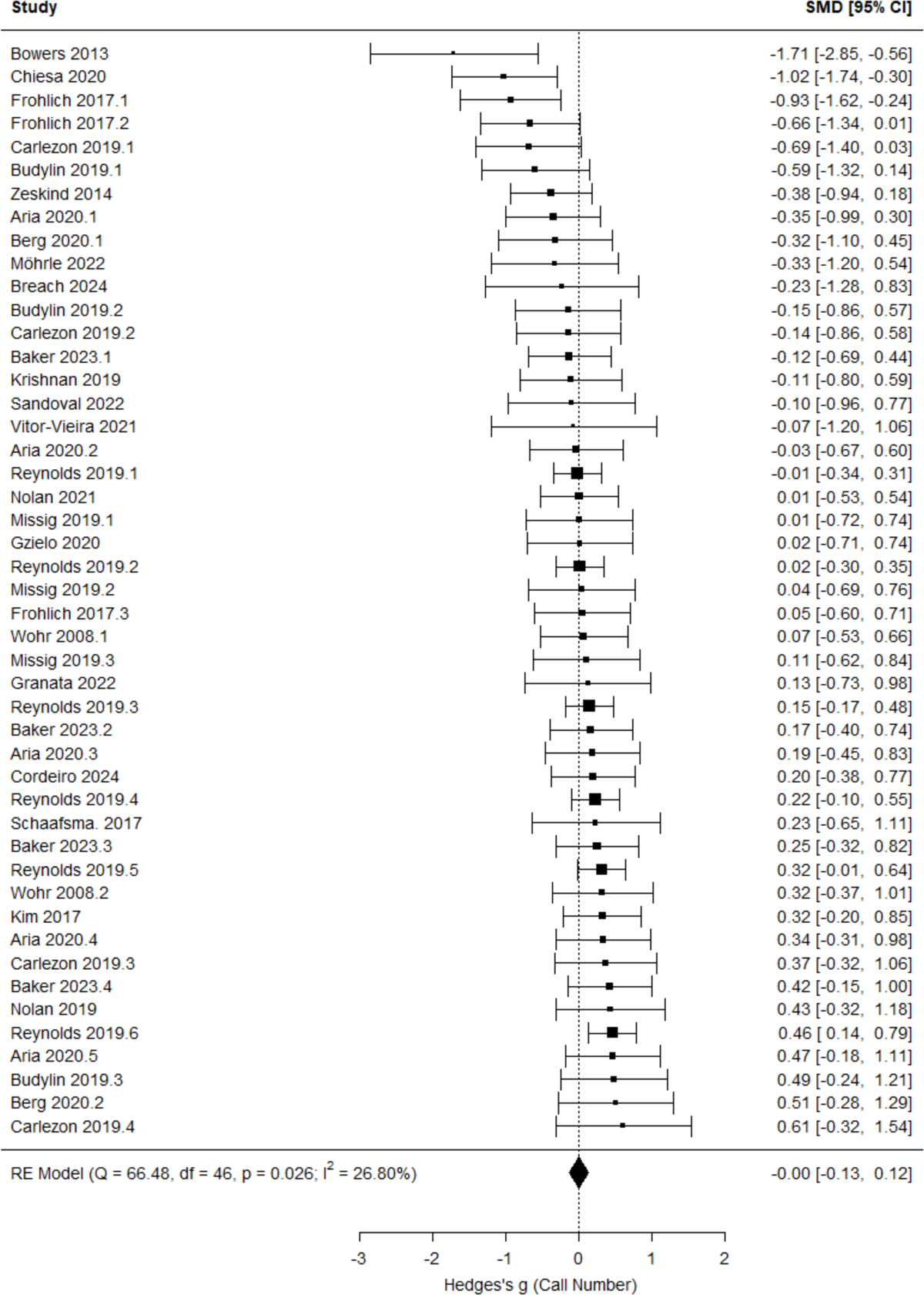
Meta-analysis results of neonatal USV call number in response to brief maternal separation.

**Supplemental Figure 2.**
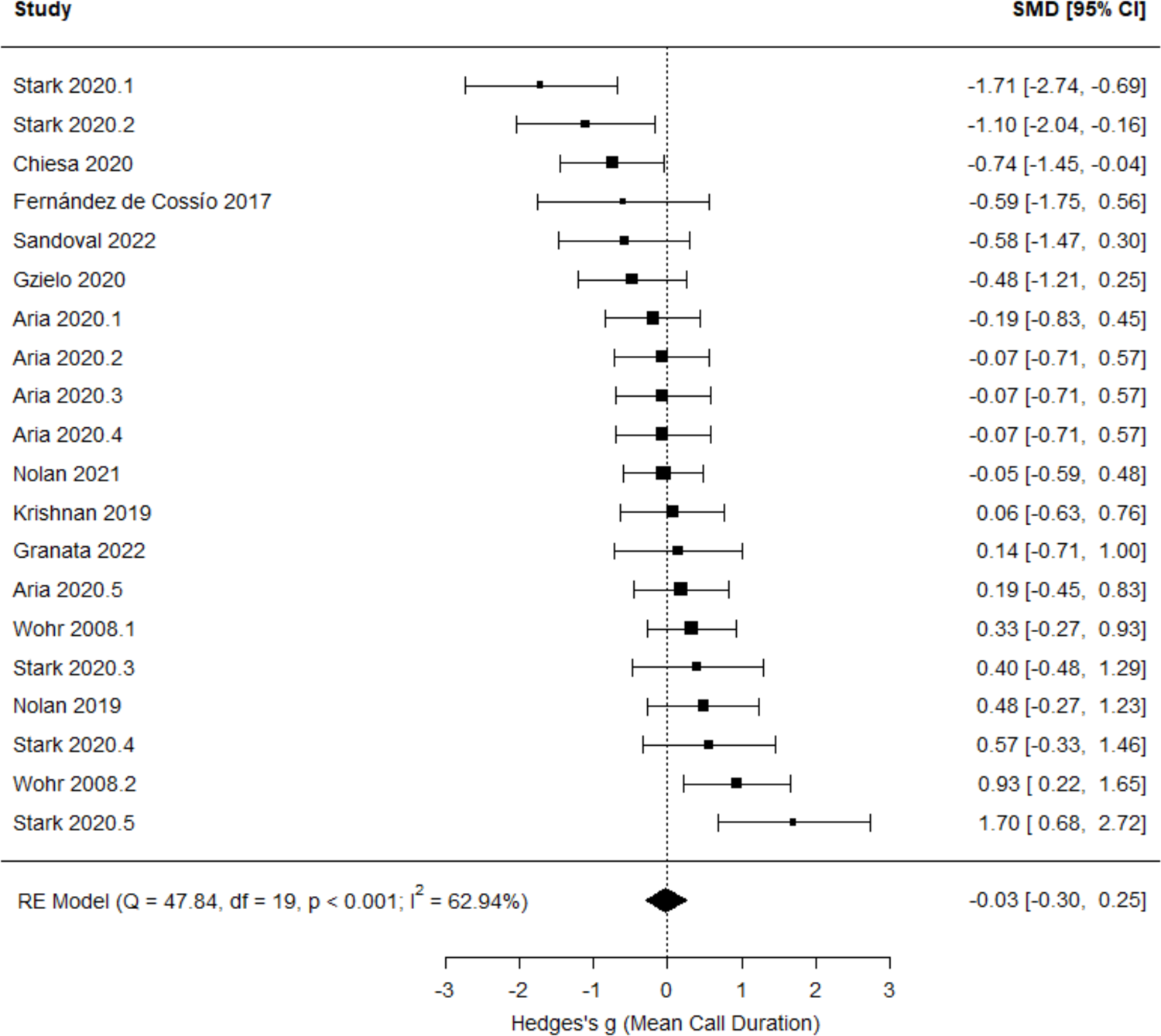
Meta-analysis results of neonatal USV mean call duration in response to brief maternal separation.

**Supplemental Figure 3.**
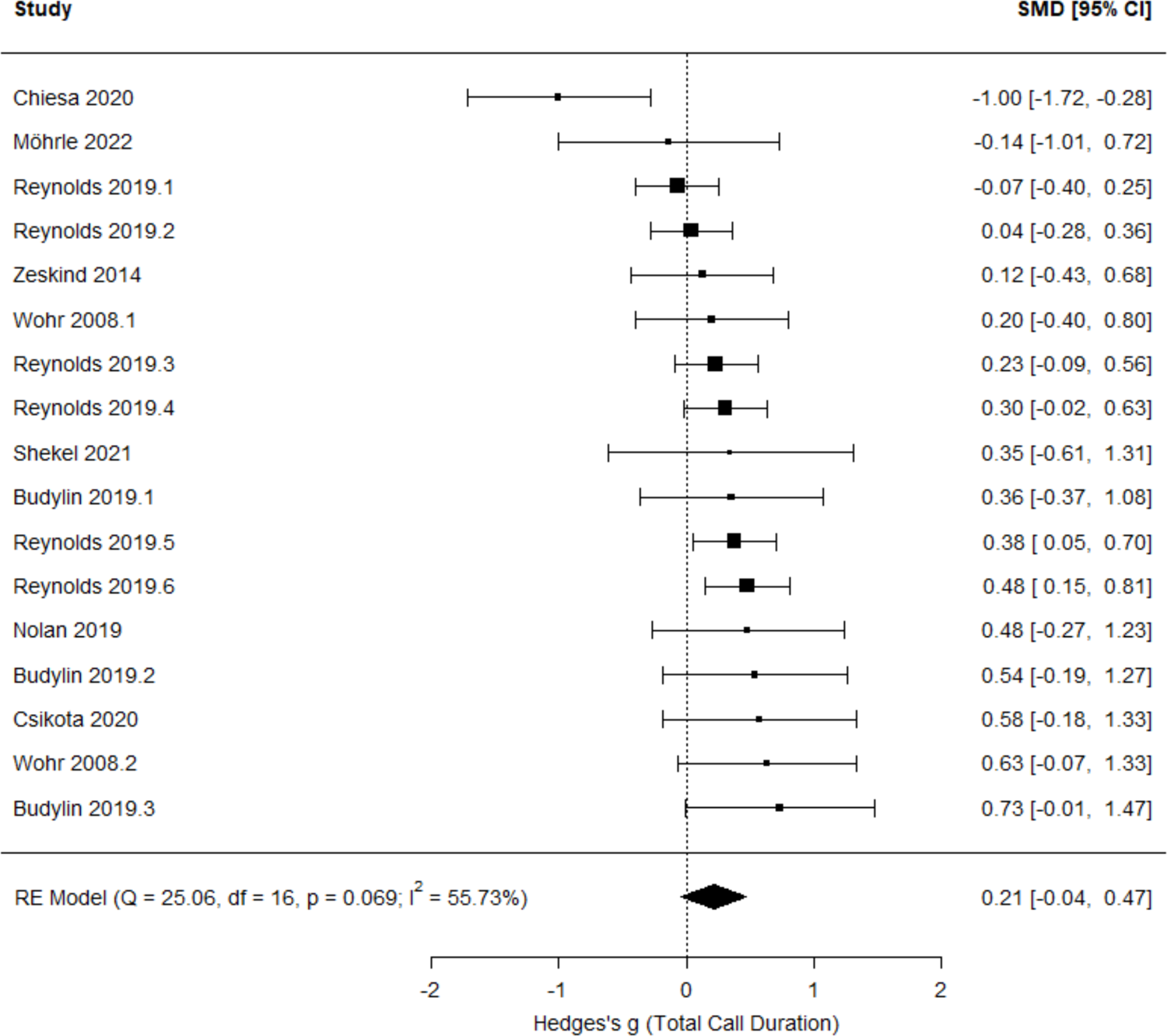
Meta-analysis results of neonatal USV total call duration in response to brief maternal separation.

**Supplemental Figure 4.**
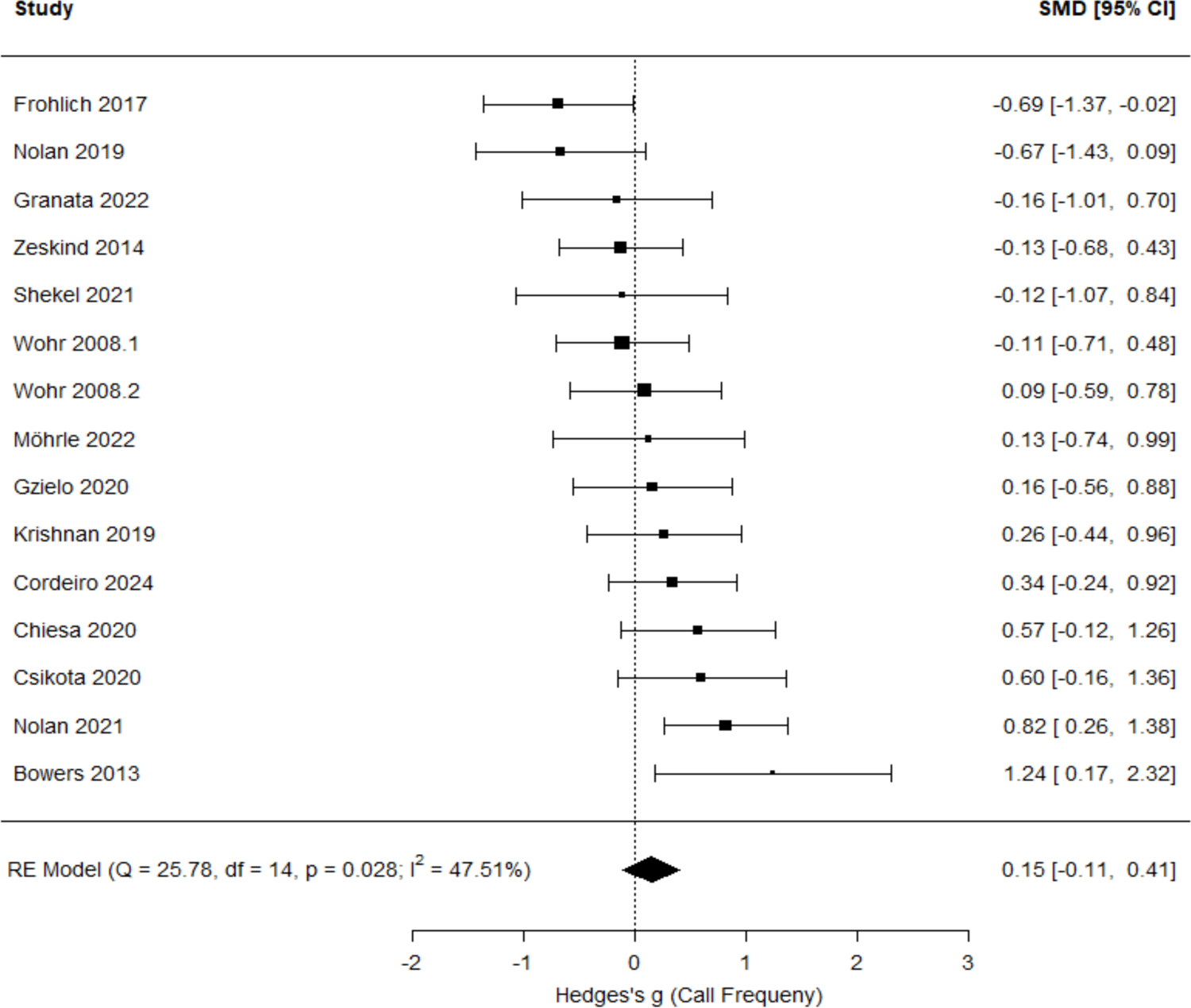
Meta-analysis results of neonatal USV call frequency in response to brief maternal separation.

**Supplemental Figure 5.**
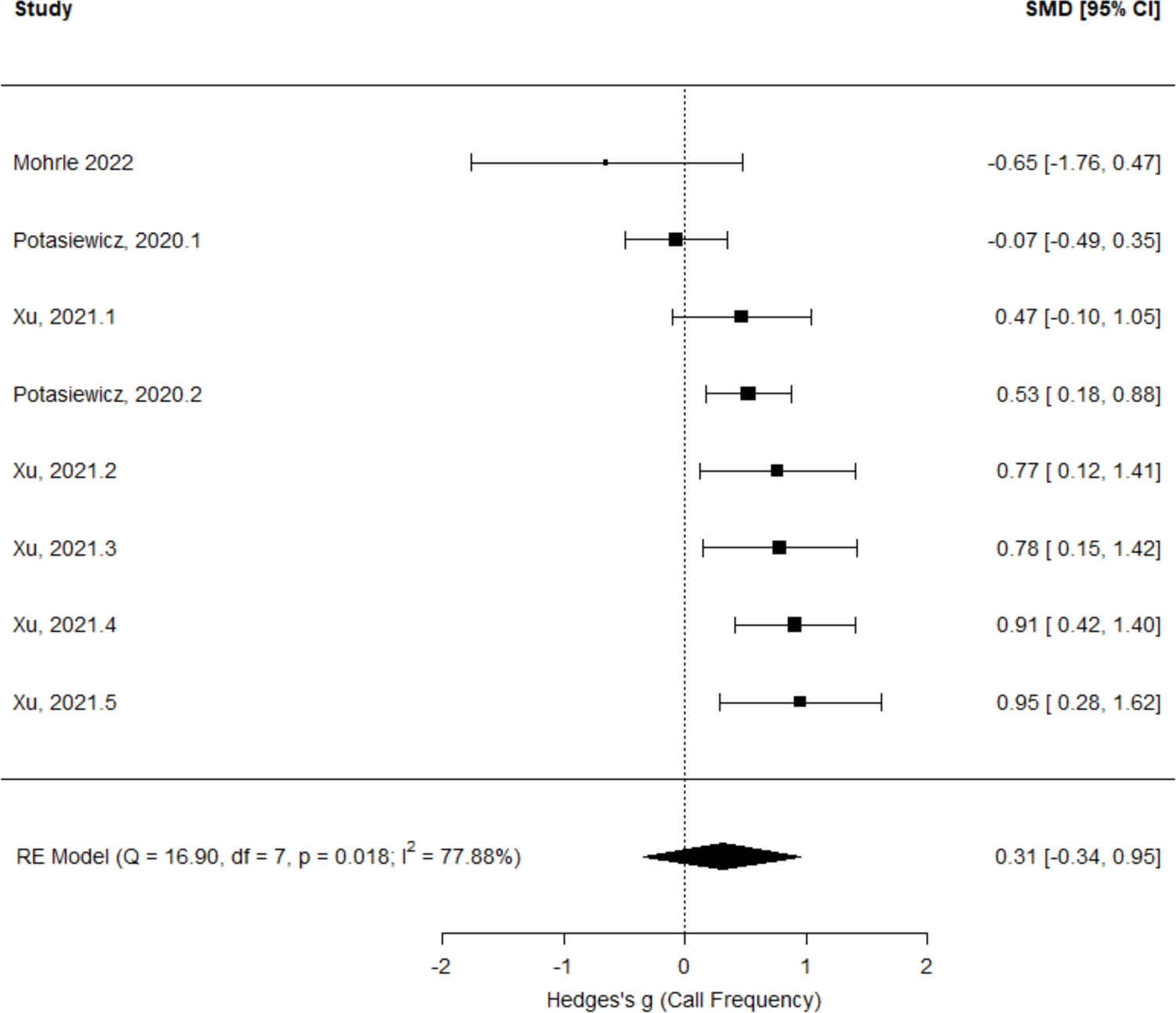
Meta-analysis results of neonatal USV call frequency difference between control and maternal immune activation male rodents

**Supplemental Figure 6.**
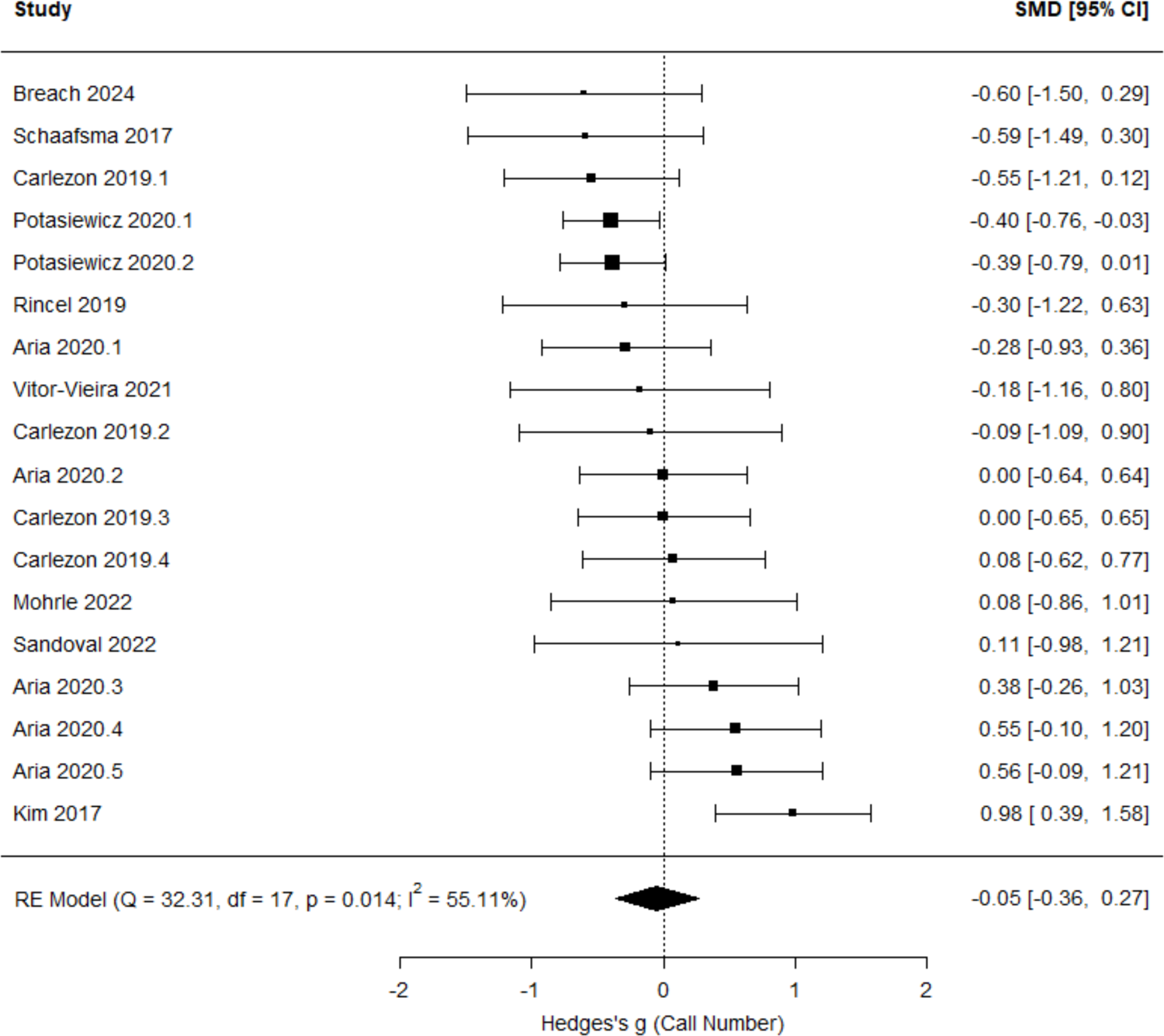
Meta-analysis results of neonatal USV call number difference between control and maternal immune activation female rodents

**Supplemental Figure 7.**
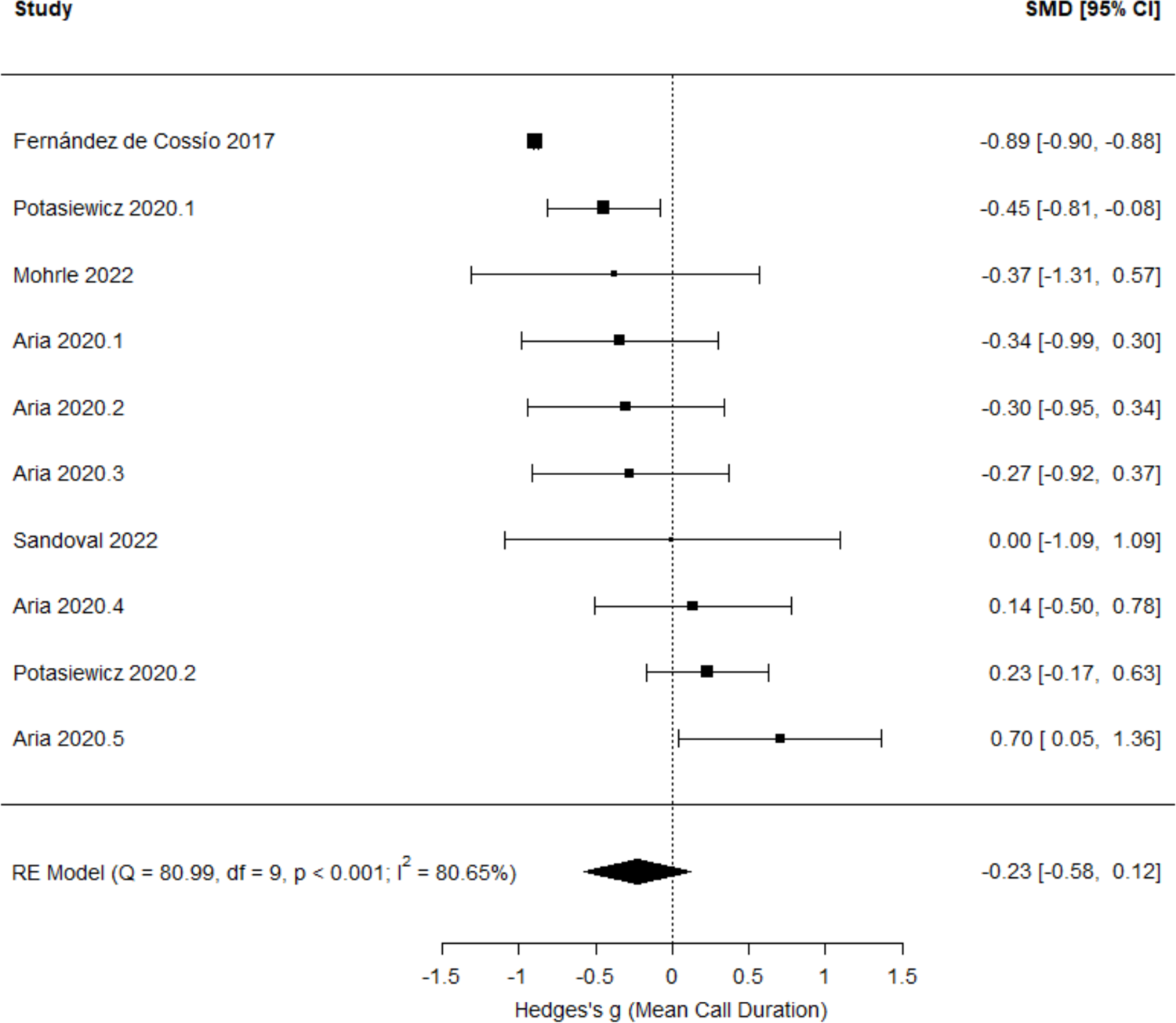
Meta-analysis results of neonatal USV mean call duration difference between control and maternal immune activation female rodents

**Supplemental Figure 8.**
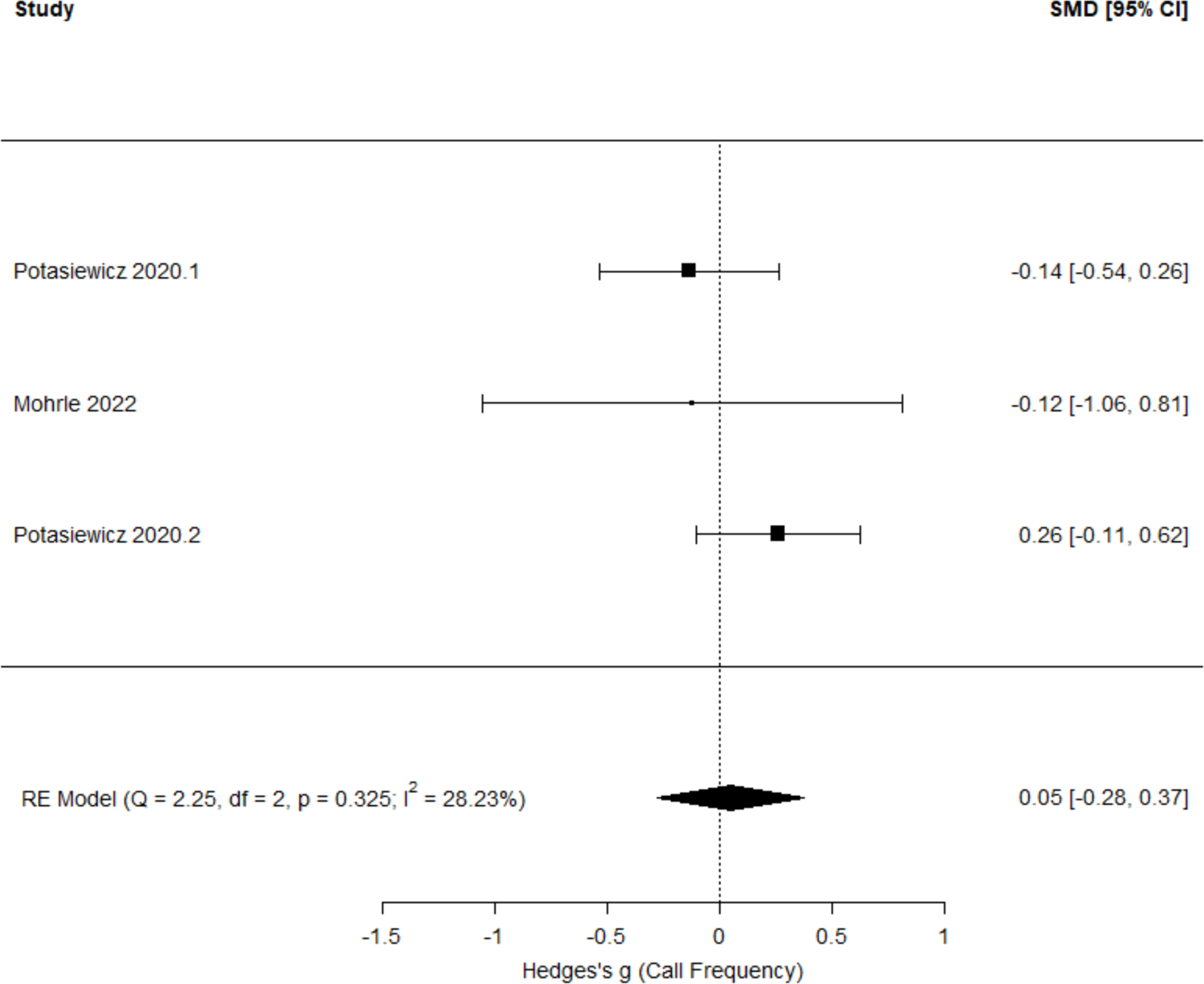
Meta-analysis results of neonatal USV call frequency difference between control and maternal immune activation female rodents

